# Genetic and environmental interactions outweigh mitonuclear coevolution for complex traits in *Drosophila*

**DOI:** 10.1101/2025.11.24.689096

**Authors:** Leah J. Darwin, Faye A. Lemieux, Rebecca Z. Bachtel, Jack H. Blocker, Camille P. Brown, Alexander D. Griffin, Jacob D. Lerman, Olivia C. Maule, Yevgeniy Raynes, David M. Rand

## Abstract

The interdependent relationship between mitochondrial and nuclear genomes is a powerful model for understanding how epistasis shapes the architecture and evolution of complex traits. Once considered a neutral marker, mitochondrial DNA variation is now recognized as critical to phenotypic evolution because of its epistatic interactions and history of coevolution with the nuclear genome. A central challenge in evolutionary genetics is to quantify the relative importance of stabilizing and directional selection shaping complex trait distributions within and among species. Both can act on interacting and/or co-evolving genes contributing to quantitative traits, but resolving their relative roles is complicated by the complex architecture of most traits. Here, we use a panel of 90 *Drosophila* mitonuclear genotypes to quantify the relative contributions of mitochondrial, nuclear, and environmental variation and their interactions to four metabolically demanding complex traits. We sample both within-species and between-species mitochondrial variation and observe stronger interaction effects attributable to within-species variation, consistent with stabilizing selection maintaining mitonuclear function. Additionally, culturing the flies on a mitochondrial Complex I inhibitor, rotenone, reveals significant genotype x environment (G×E and G×G×E) interaction effects, providing insight into how genetic variation can be maintained across changing environments. Our results have broader implications in medicine, where mitochondrial DNA donors with longer purifying selection histories may be safer for mitochondrial replacement therapies.

## Introduction

The observation that many traits vary continuously among individuals raises fundamental questions about the genetic basis and maintenance of phenotypic diversity. These quantitative traits are often challenging to study because they arise from the combined segregation of many polymorphic loci and are sensitive to different environmental contexts [1]. Interactions between genes (G×G, epistasis) and between genes and the environment (G×E) can further complicate trait expression, leading to unpredictable outcomes. The extent to which quantitative traits are shaped by additive versus interactive effects is a matter of considerable debate. Some argue that although gene interactions contribute to variation in most biological systems, population-level genetic variance is mostly additive [2]. Others contend that epistasis is pervasive and that additivity can arise as an emergent property of complex epistatic networks [3]. Interaction effects could help to explain “missing” heritability common in genome-wide association studies [4]. They may also account for how genetic variation can be maintained under stabilizing selection [5, 6], which appears to be a major force acting on quantitative traits [7].

Fully comprehensive tests for epistatic effects on traits are both experimentally and computationally intractable because the number of variant combinations– and thus statistical model terms or experimental treatments– increases exponentially as more loci are considered. One effective strategy to address this challenge is to reduce the number of testable interactions by considering sets of genes rather than individual gene combinations. A natural example of such sets is the mitochondrial (mtDNA) and nuclear genomes, which jointly encode oxidative phosphorylation (OXPHOS) complexes responsible for energy synthesis [8]. Their interdependence and the functional importance of OXPHOS to most traits makes the mitonuclear system a powerful model for studying adaptive evolution [9, 10]. Much work has focused on the adaptive divergence of mitochondrial genes and the response of the nuclear genome [11, 12], with the expectation of positive selection on nuclear genes compensating for deleterious mutations in mtDNA. This model predicts that mismatched mitochondrial and nuclear genomes would reduce hybrid fitness, observed in systems such as marine copepods [13], yeast [14], fruit flies [15], nematodes [16], and swordtail fishes [17]. However, analyses of the sites of amino acid contact between mtDNA- and nuclear-encoded proteins in bivalves [18] and mammals [19] have not revealed consistently significant dN/dS ratios greater than 1, which would be expected under positive selection. Some instances of elevated dN/dS for these proteins are attributed to relaxed rather than positive selection [20] suggesting that such incompatibilities may be the exception rather than the rule [21]. Conflicting empirical findings highlight the need for alternative models of mitonuclear coevolution and its impact on quantitative traits.

In this study, we constructed a panel of mismatched mitonuclear genotypes in *Drosophila* to elucidate the contributions of genetic and environmental interaction effects to four metabolically demanding traits: body weight, climbing velocity, flight performance, and development time. We introduced 10 mtDNAs from each of two diverged natural populations of *D. melanogaster* and two additional species (*D. simulans* and *D. yakuba*) onto two common *D. melanogaster* nuclear backgrounds. To assess how genetic effects and interactions are influenced by environmental perturbation, each line was exposed to the OXPHOS Complex I inhibitor rotenone. Rotenone is a naturally occurring plant compound that is widely used as a pesticide to control insects, mites, ticks, and undesirable fish in a variety of environmental contexts [22]. It is also used to induce Parkinson’s disease phenotypes in model organisms due to its targeted effect on mitochondria in specific neurons. We chose rotenone as an environmental variable based on its targeted inhibition of Complex I, a protein complex that is encoded by both mitochondrial and nuclear genes, allowing a direct manipulation of the mitonuclear genetic architecture of interest.

By quantifying how G×G and G×E interactions shape phenotypic variation within populations, our results show how mildly deleterious mtDNA mutations that persist under stabilizing selection influence trait variation. Understanding the impact of mildly deleterious mutations in different genomic and environmental contexts provides a framework for anticipating their consequences in clinical settings such as mitochondrial replacement therapy [23, 24]. Our results further demonstrate that purifying selection acting on mtDNA prevents the accumulation of deleterious alleles, challenging compensatory models of mitonuclear coevolution.

## Results

In this study, we constructed a panel of 80 intraspecific and eight interspecific mitonuclear genotypes. These comprised 22 mitochondrial haplotypes, including within-species variation from Beijing and Zimbabwe *D. melanogaster* (*D. mel*) populations and from two related species, *D. yakuba* (*D. yak*) and *D. simulans* (*D. sim)* (previously placed on *D. mel* nuclear backgrounds; see Methods). Virgin females carrying each mitochondrial haplotype were backcrossed for 10 generations to males of two North American *D*.*mel* nuclear backgrounds, OreR and DGRP375 (hereafter labeled as mtDNA;nucDNA, e.g., Bei05;OreR or ZW144;375). This process of backcrossing was done in duplicate, creating two “builds” of each mitonuclear line to control for potential nuclear variation (theoretically 1/2^10^, or 0.0976% of the nuclear genome after 10 generations of backcrosses) that could confound the effect of mtDNA genotype on phenotype. We report significant correlations between builds for all traits (Supplementary Fig. 1). An additional two lines retaining their “native” mitochondrial haplotypes, OreR;OreR and DGRP375;DGRP375 (Figure 1a) were included, totaling 90 assayed lines over the course of this study.

**Figure 1:**
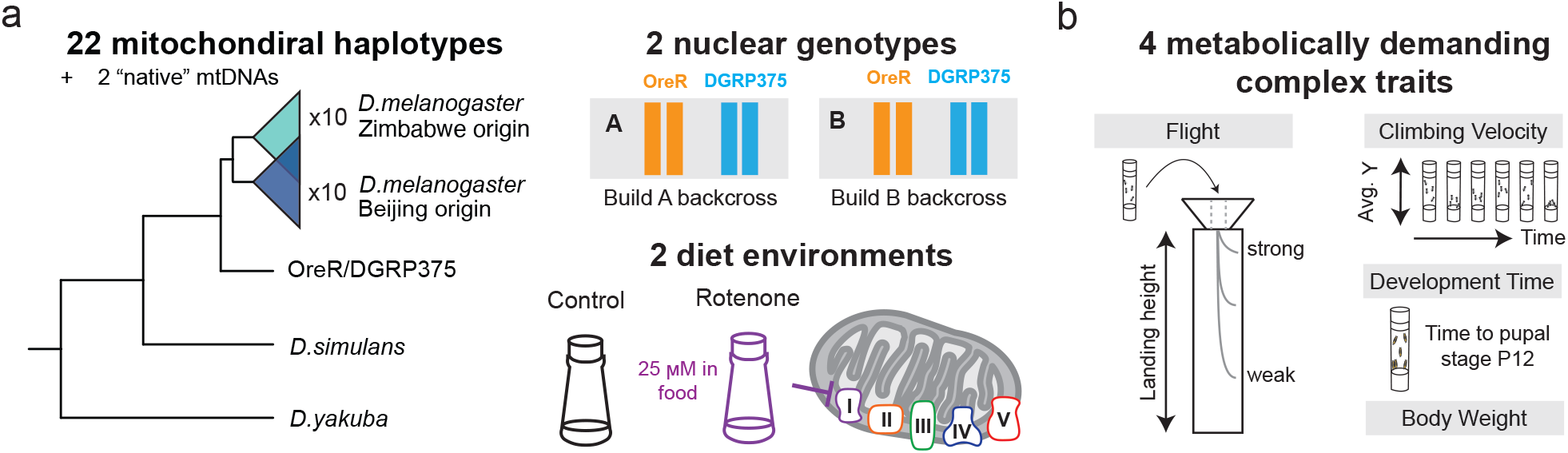
Experimental design and genotypes sampled. **a** Mitochondrial phylogeny shows 10 haplotypes from a *D. melanogaster* population in Zimbabwe, 10 from Beijing, 2 from *D. simulans* and *D. yakuba*, and the two “native” mtDNAs corresponding to the two nuclear genotypes used. Each mtDNA was backcrossed in duplicate builds A and B to the OreR and DGRP375 North American *D. melanogaster* nuclear backgrounds and exposed to control and rotenone diets. Rotenone is a mitochondrial Complex I inhibitor and was administered at 25*µ*M. **b** Four quantitative traits were measured: flight performance, climbing velocity, body weight, and development time. Flight performance was assessed by measuring the landing height of individuals when dropped. Climbing velocity was measured by regressing average (avg.) y-position against time for a replicate vial of flies. In total, 90 total lines were quantified across nine experimental blocks (2,160 assays).

We measured the quantitative traits development time, climbing velocity, flight performance, and body weight in two environments: a control environment and one containing rotenone, a naturally occurring OX-PHOS Complex I inhibitor, at a sublethal dose (Figure 1b). For the weight, climbing, and flight traits, males and females were sorted and assayed separately. There were no significant correlation coefficients between any combination of two phenotypes for both females and males (Supplementary Fig. 2, Supplementary Tables 1–2). We assessed the reproducibility of our trait measurements by retesting a subset of lines. For all retested phenotypes, there were significant rank correlations between the original measurements and the repeated ones (*p* < 0.01, Supplementary Fig. 3, Supplementary Table 3).

### Nuclear, environmental, and mitochondrial main effects and interactions

We used a mixed-effect linear model to quantify the effects of the nuclear genotype, rotenone treatment, mitochondrial genotype and their interactions on phenotype (Equations 1–2). Results for the climbing, flight, and weight traits focus on a sex-stratified model (Equation 2) with a Bonferroni corrected significance threshold (*α* = 0.05/4 traits = 0.0125) and used a semi-partial 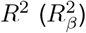 to quantify relative effect sizes. Across all phenotypes, the nuclear main effect was significant in both sexes (*p* < 0.001)(Figure 2a, Supplementary Tables 7–9). In both males and females, the nuclear genome had the largest effect for climbing (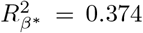 for females; 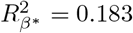 for males), flight (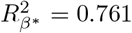 for females; 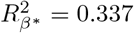 for males), and development time (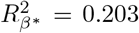, not stratified by sex) (Supplementary Tables 7–9). We found that animals with the DGRP375 nuclear background developed slower than those with the OreR background but performed better in flight and climbing, which may be related smaller body weight (Figure 2a). Additionally, principal component (PC) analysis of all phenotypes showed that the nuclear genome had the largest effect on variance, as PC1 (reflecting nuclear background effects) explained 51.11% of the total variance (Supplementary Fig. 4). The effect of rotenone was significant for the climbing trait (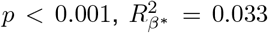 for females; *p* < 0.001, 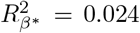 for males), development time (*p* < 0.001, 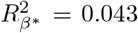), and flight in males (*p* < 0.001, 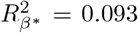) (Figure 2b, Supplementary Tables 7–9). Treatment with rotenone negatively affected most traits, reducing flight and climbing performance and slowing development time (Figure 2b). Notably, we find that survivorship bias was unlikely to affect our overall estimations of variance. When a subset of 10 lines were tested for differences in survival on rotenone diets (see Methods), there was no significant treatment effect for survival from egg to adult (*p* = 0.59, Supplementary Table 4). Further, the estimated confidence intervals for total surviving adults for each diet were overlapping (Supplementary Table 5), indicating a negligible treatment effect on survival. Additionally, females did not preferentially lay eggs on either food type (*p* = 0.086, Supplementary Table 6).

**Figure 2:**
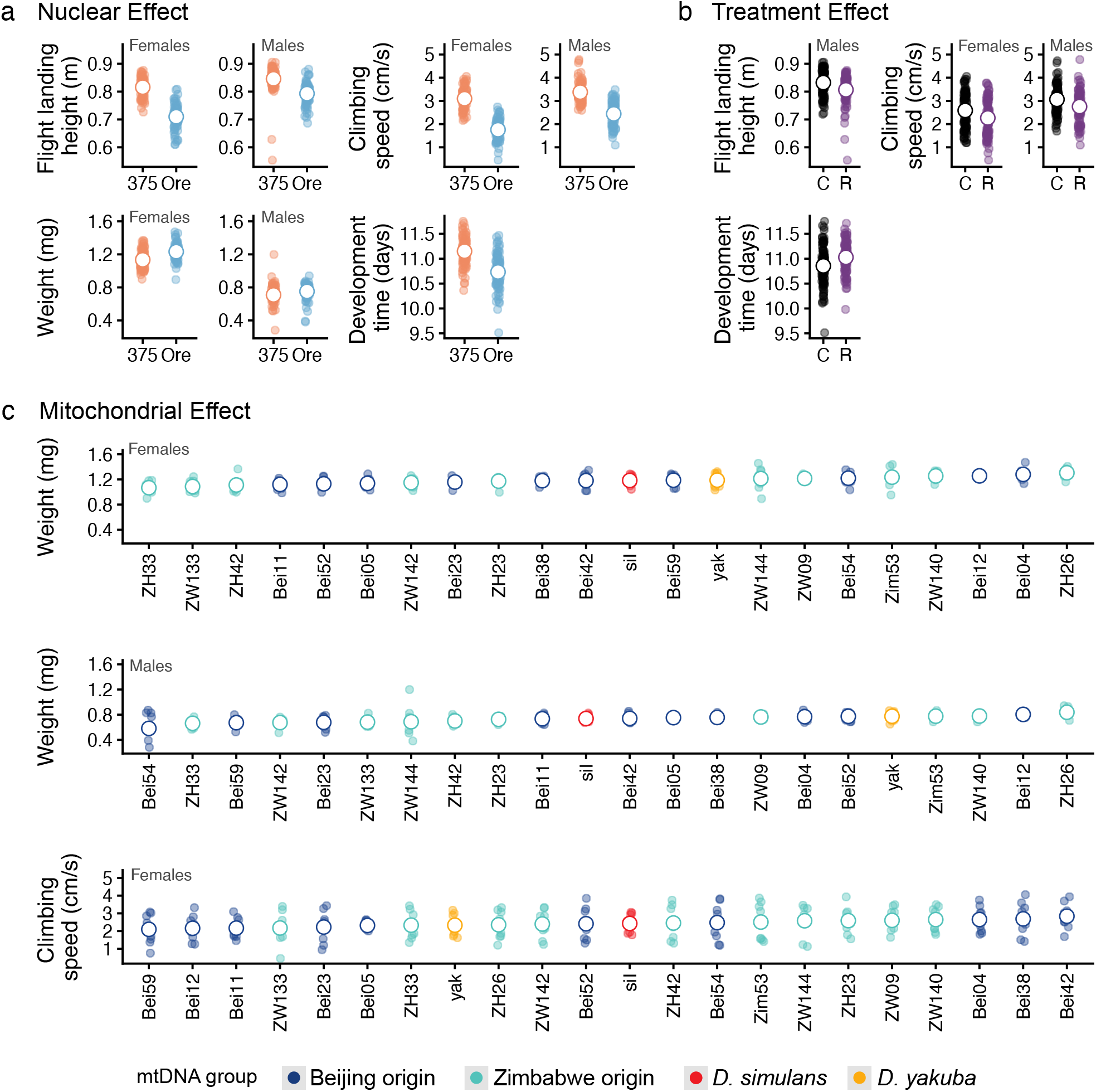
Significant main effects. For each tested effect, trait/sex combinations significant at a Bonferroni corrected significance threshold *α* = 0.05/4 traits = 0.00125 are shown. Data point clouds show the average adjusted trait values for all 88 mitonuclear genotypes. Note trait/sex combinations that are not shown were quantified but not statistically significant (*p* > 0.00125). **a** Nuclear effects across the DGRP375 (375) and OreR (Ore) genotypes. The large white circle is the estimated marginal mean of each nuclear background. **B** Treatment effects across the control (C) and rotenone (R) environments. The large white circle is the estimated marginal mean of each treatment. **c** Mitochondrial effects across the Beijing, Zimbabwe, *D. simulans* (siI) and *D. yak* (yak) groups. The large white circle is the estimated marginal mean of each mtDNA.

The mitochondrial genome had significant main effects for weight (*p* < 0.001, both sexes) and for female climbing (*p* = 0.005) (Figure 2c, Supplementary Tables 7–8). For the weight phenotype, the mitochondrial effect size was particularly notable as it was greater than the nuclear effect in both females 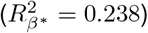, and males 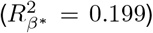 (Supplementary Tables 7–8). The significant mtDNA effects were not driven by similar effects from within either the Beijing or Zimbabwe *D. mel* populations, or by extreme phenotypes from the two outgroup species *D. sim* and *D. yak*. Rather, when the estimated marginal means of the mitochondrial effects were ranked, the Beijing and Zimbabwe mtDNAs are not distinctly clustered and the *D. sim* and *D. yak* mtDNAs were in the middle of this range (Figure 2c). To further investigate the relationship between the mitochondrial phylogeny and phenotypes across the four traits within nuclear background, sex, and treatment groups, we measured phylogenetic signal. Specifically, we used Blomberg’s *K*, which tests for a relationship between a phylogeny and a trait. Under a mitonuclear coadaptation model, divergent mtDNAs (*D. yak, D. sim*) in a *D. mel* nuclear genomic background should disrupt mitonuclear communication and have larger phenotypic effects than conspecific mtDNAs. Conversely, variation within *D. mel* populations (Beijing vs. Zimbabwe) should show greater similarity within clades if traits have been shaped by shared selective pressures. Our tests revealed no significant Blomberg’s *K* values for all combinations of sex, treatment, nuclear background, and phenotype (Supplementary Table 10). Instead, the outgroups had intermediate trait measurements relative to the Beijing and Zimbabwe clades for all traits (control females shown in Supplementary Fig. 5).

Like the main effects, second-order interaction variance also differed across traits and sexes, indicating that each trait had a distinct architecture. The mitonuclear genetic epistasis (G×G) was significant only for male weight (*p* < 0.001) (Figure 3a, Supplementary Table 8). In the climbing trait, the G×G interaction was significant at a less conservative threshold *α* = 0.025 in males (*p* = 0.015) and marginally insignificant in females (*p* = 0.026) (Supplementary Tables 7–8). We found the mitochondrial-by-treatment G×E to be more common across traits. The mtDNA G E effect was significant in the development time phenotype (*p* = 0.0123) and in both females (*p* < 0.001) and males (*p* < 0.001) in the climbing phenotype (Figure 3b, Supplementary Tables 7–9). In the development time trait, the mitochondrial-by-treatment interaction effect size 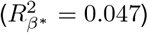 was slightly larger than the treatment main effect size 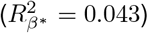 (Supplementary Table 9). For all significant second order interaction effects, the slopes of the reaction norms were not more similar within mtDNA group (Figure 3). Furthermore, the two outgroups *D. sim* and *D. yak* appeared to be less reactive than many of the *D. mel* mtDNAs (Figure 3).

Interaction effects can be highly sensitive to the specific mtDNA genotypes included in the study relative to main effects because they are measured through comparisons of comparisons. It may be the case that interactions are being driven by a few specific mtDNA genotypes rather than a general “stable” feature of the data. To test the stability of these reported effects, we refit the model on 1000 unique combinations without replacement for subsets of size *k* = (5, 10, 15, 20) mtDNAs (for *k* = 20, all 231 combinations are retested). The mitonuclear genetic epistasis (G×G) in the male weight trait was highly unstable across subsets of the data (significant in 8%, 11%, 16% and 7% of retests for *k* = (5, 10, 15, 20), respectively, Supplementary Figure 6).

Significant *p*-values were outliers, indicating that this interaction was driven by a few mtDNAs rather than a general property of the entire panel. We additionally found that weight in both sexes was negatively related to estimates of larval density (Supplementary Figure 7), therefore the interaction effect may instead be capturing environmental confounding that is unevenly distributed across lines. Importantly, the G×E interaction effects were more stable across subsets of the data. For development time, the number of significant subsets increased with the size of *k* (significant in 11%, 16%, 26% and 43% of retests for *k* = (5, 10, 15, 20), respectively, Supplementary Figure 6) suggesting that the interaction effect is distributed across the genotypes and detectable as more genotypes are sampled. For climbing speed, the G×E interaction effect was highly stable across all levels of *k* for both sexes (significant in 74%, 98%, 100% and 100% of retests for females; and 70%, 98%, 100% and 100% for males for *k* = (5, 10, 15, 20), respectively, Supplementary Figure 6) indicating that this interaction is a general property of the climbing trait.

**Figure 3:**
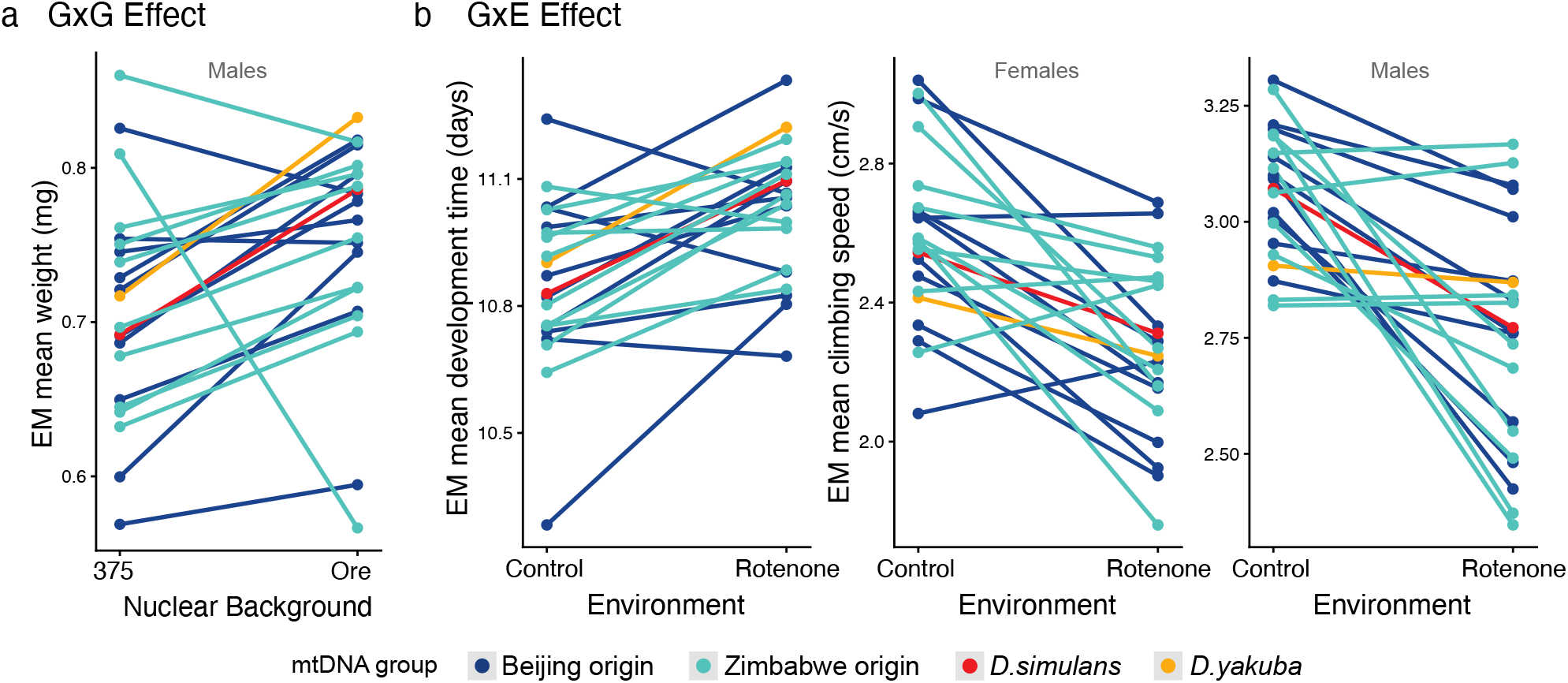
Significant second order interactions. For each tested effect, trait/sex combinations significant at a Bonferroni corrected significance threshold *α* = 0.05/4 traits = 0.00125 are shown. Note trait/sex combinations that are not shown were quantified but not statistically significant (*p*>0.00125). Lines crossing indicate an interaction. **a** Mitonuclear GxG effects shown by the estimated marginal (EM) mean for the male weight trait across the two nuclear background for each mtDNA. **b** Mito-rotenone GxE effects for development time and female and male climbing. EM means for trait are given across the control and rotenone environments.

### Mitochondrial genotype performance across nuclear and environmental contexts

We identified a significant third order G×G×E interaction effect in the climbing trait in both females (*p* < 0.001, Supplementary Table 7) and males (*p* < 0.001, Supplementary Table 8). Notably, this higher order interaction effect was very robust across resamples of subsets of *k* mtDNAs, as similarly tested for second order interactions (significant in 53%, 80%, 96% and 100% of retests for females; and 87%, 100%, 100% and 100% for males for *k* = (5, 10, 15, 20), respectively, Supplementary Figure 6). To further explore how mitochondrial genotypes differ phenotypically across nuclear genotype and treatment combinations, we used an additive main effects and multiplicative interaction (AMMI) model (Equation 3). This method applies principal component analysis (PCA) to decompose interaction effects after centering on additive effects, visualized with a biplot of genotypes and environments. We combined nuclear background and treatment environment into a single composite “environment” for the mitochondrial term, as these factors jointly constitute the context within which mitochondrial genotype effects are expressed. This allows the biplot to visualize third-order interaction effects (Mito:Nuc:Treatment) while also absorbing second-order effects (Mito:Nuc and Mito:Treatment), though some interaction structure may be lost through this condensation [27]. Points (here, mtDNAs) near the interaction PCA (IPCA) axes origin are more stable, whereas those farther away show stronger interactions [28]. Acute angles between genotype points and environment vectors indicate positive interactions (e.g. ZW144 and O:C in Figure 4a inset), and obtuse angles (e.g. ZW144 and O:R in Figure 4a inset) indicate negative ones. Figure 4 shows the AMMI biplot of the first two IPCA axes for the climbing trait. IPC1 was significant in females (46.2%, *p* < 0.0001) and males (58%, *p* < 0.0001), IPC2 was also significant (37.2%, *p* = 0.004 for females; 37.8%, *p* = 0.004 for males), IPCA3 was not significant in either sex and is excluded.

**Figure 4:**
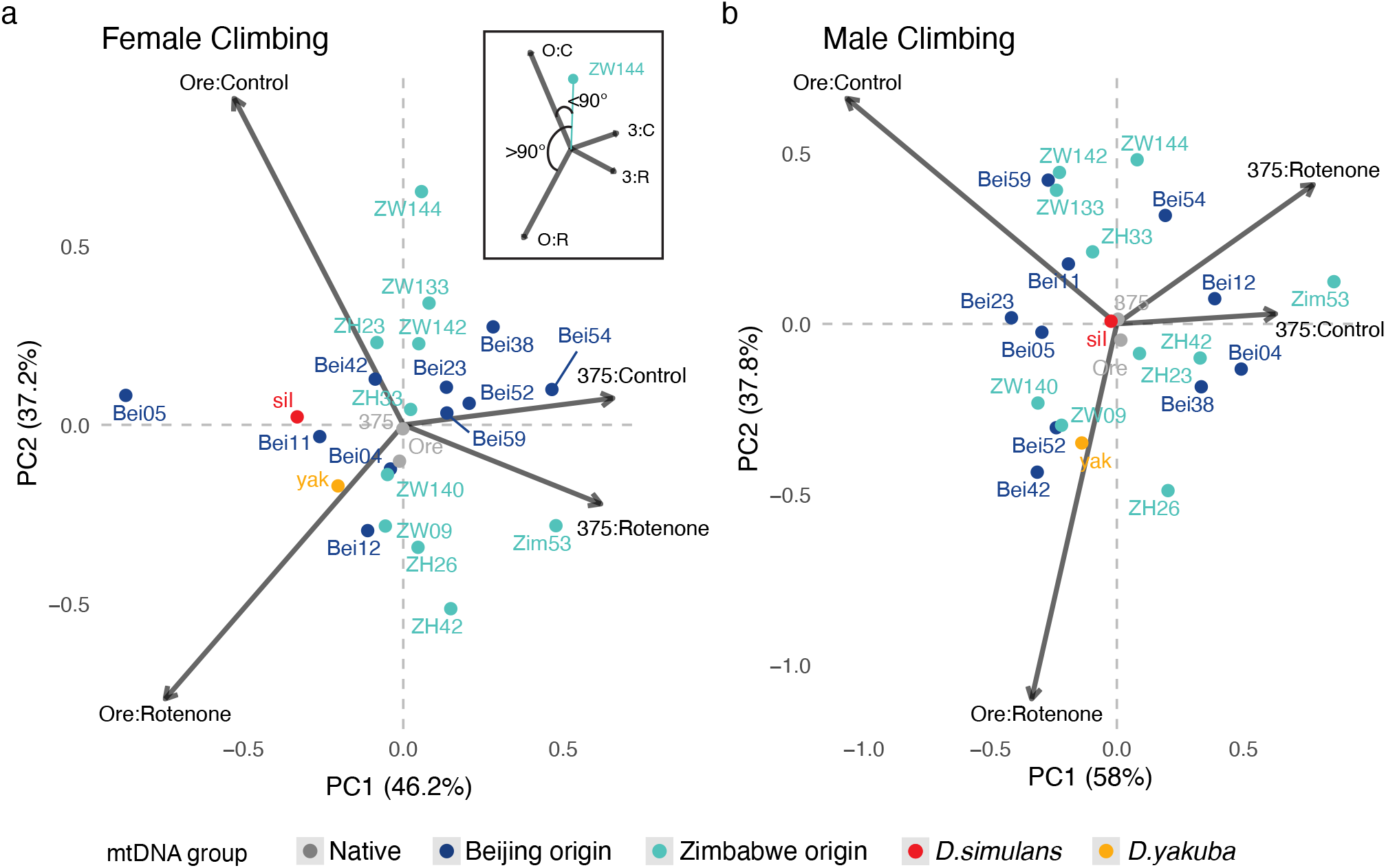
AMMI biplot of first two interaction PCs for the climbing trait. Nuclear:Treatment combinations are represented by arrows. Mitochondrial genotypes are represented by points and colored by general grouping (Beijing origin, Zimbabwe origin, *D. simulans, D. yakuba*, and the native mitochondria 375/Ore.) Mitochondrial genotypes closer to the origin show smaller interaction effects and are generally stable across different Nuclear:Treatment combinations. Genotypes further from the origin show greater interaction effects and align closer (acute angle, ZW144 and O:C in inset A) to environments where there is a positive interaction and are more distant (obtuse angle, ZW144 and O:R in inset A) where there is a negative interaction. Data from female individuals is shown in panel (a) and males in panel (b).

Using the AMMI model, we observed that OreR nuclear background causes stronger climbing performance interactions among mitochondrial genotypes than the DGRP375 background. In both females (Figure 4a) and males (Figure 4b), the OreR nuclear background environment on both control (Ore:Control arrow) and rotenone (Ore:Rotenone arrow) food are much further from the origin than the DGRP375 nuclear backgrounds (375:Control and 375:Rotenone arrows) indicating stronger interaction effects.

There is no trend that suggests that certain mitochondrial groups (i.e. haplotypes from within Beijing or Zimbabwe) performed similarly better or worse on any given environment. However, there were some haplotypes from within the Beijing and Zimbabwe groups that stood out as having large interaction effects. In both males and females, ZW144 displayed a strong interaction effect indicated by its distance from the origin (Figure 4). We also observed that in both sexes, the “native” mitochondrial haplotypes 375 and Ore were close to the origin indicating these were stable across the treatment environments. These haplotypes were not reciprocally placed onto the two nuclear backgrounds so consequently cannot deviate along the horizontal plane.

Surprisingly, the interaction effects for both *D. sim* and *D. yak* were not stronger than those observed within either *D. mel* group despite having diverged as species ∼2.5M and ∼10M years ago, respectively. This is apparent from the placement of the *D. sim* and *D. yak* points inside the ellipse of points defined by either the Beijing or Zimbabwe mtDNAs from *D. mel*. Interestingly, the strength of interaction effects appears to be unrelated to amino acid (AA) divergence between mtDNA haplotypes: relative to the *D. mel* mtDNA reference genome, there are 142 changes between *D. mel* and *D. yak*, 92 between *D. mel* and the *D. sim* siI, 12 across all *D. mel* Beijing mtDNAs, and 11 across all *D. mel* Zimbabwe mtDNAs (Supplementary Table 11). We then considered whether mitonuclear interactions could be attributable to AA changes at physical protein-protein contact sites between mtDNA and nucDNA residues, since OXPHOS Complex I (CI) inhibited by rotenone comprises both mitochondrial and nuclear encoded proteins. We used a Cryo-EM structure of CI [29] to map the missense changes to contact and non-contact positions and found that 62.5% of AA mutations among the 20 *D. mel* mtDNAs were in CI nuclear contact sites compared to 31.3% and 33.6% of the AA mutations fixed between *D. mel* and *D. sim* or *D. yak*, respectively (Figure 5, Supplementary Table 12). To assess the impact of amino acid changes, we classified them as conservative or radical, with radical changes involving changes between AA classes (nonpolar or uncharged polar) [30]. A higher proportion of radical substitutions occurred in *D. mel* mtDNAs (8/20, 40%) than in divergence to *D. sim* (24/92, 26%) or *D. yak* (43/142, 30%) (Supplementary Table 11). While not significant by G-tests of heterogeneity, the trend is towards more deleterious conditions within than between species.

**Figure 5:**
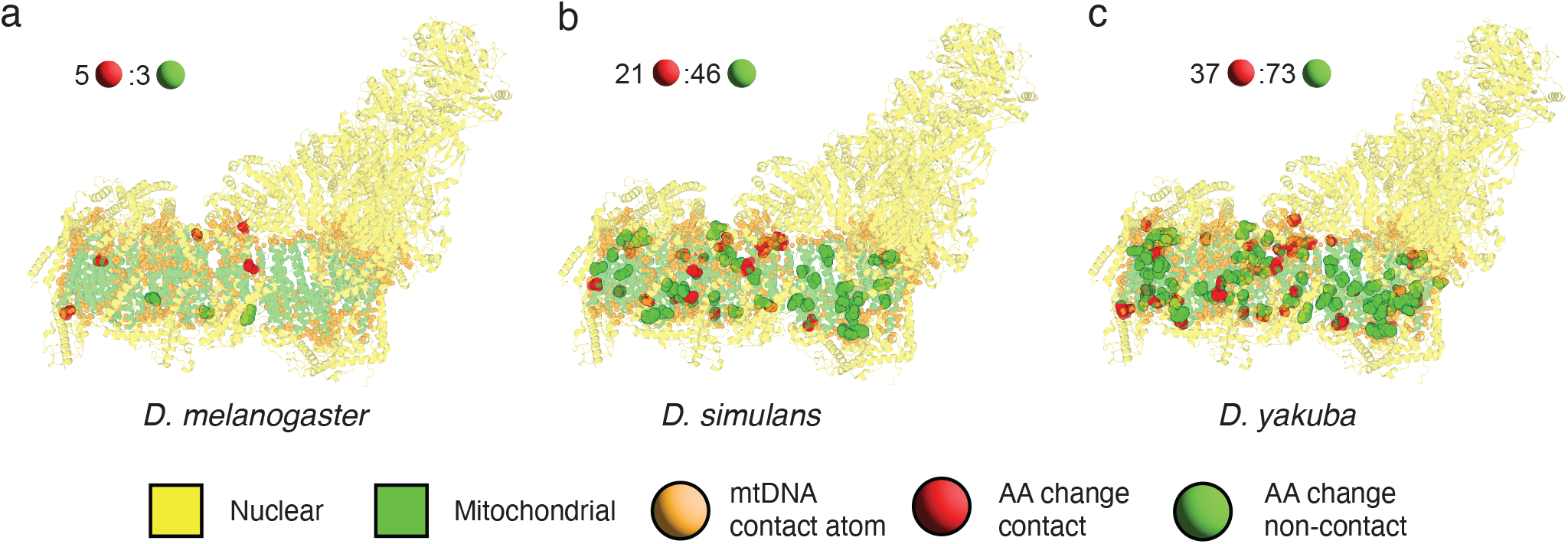
Complex I mitonuclear contact sites. Complex I (CI) protein structure (from [29]) showing both mitochondrial and nuclear encoded proteins. Atoms from mtDNA encoded proteins <4Å from nuclear residues are shown as orange spheres (contact sites). The larger green and red spheres show non-contact and contact sites respectively for each CI missense mutation in the **a** *D. melanogaster* haplotypes, **b** *D. simulans* haplotype, and **c** *D. yakuba* haplotype. Ratios of contact to non-contact AA changes for each species are given. Note that the mutations shown are not strictly private to each group and many overlap. All AA changes are given in Supplementary Table 11.

### Sex differences influence traits, but Mother’s Curse is not evident

Using the ANOVA model that included sex as both a main and interaction effect (Equation 1), we found sex to be a significant main effect in all traits (*p* < 0.001, Supplementary Table 9), with the largest effect on weight 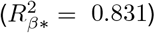 reflecting the known sexual dimorphism of body size. We also found a number of significant genotype-by-sex, treatment-by-sex and third order interactions between mitochondrial genotype, sex, and nuclear genotype (Supplementary Table 9).

We also examined how maternal inheritance has shaped mitochondrial haplotype variation. Maternal inheritance allows selection on female traits but can let male-harming mutations accumulate, a hypothesis known as Mother’s Curse [31, 32]. The weak form of this hypothesis predicts greater phenotypic variation among mtDNA genotypes in males than in females which we tested for by comparing coefficient of variation (CV) among mtDNAs in the two sexes. The strong form predicts variation in both sexes with negatively correlated male and female performance [33]. Across all 12 trait, environment, and nuclear genotype combinations there was no evidence of either weak or strong Mother’s Curse, as no CV comparisons were significant (Supplementary Fig. 8, Supplementary Table 14), and none of the significantly correlated traits showed a negative correlation (Supplementary Table 14).

## Discussion

In this study, we leveraged a genetically diverse panel of mitonuclear genotypes of *Drosophila* to evaluate the impact of mitonuclear coevolution, epistasis, and GxE on four complex traits: climbing velocity, flight performance, body weight, and development time. Unlike many systems where a full-factorial test of epistasis is infeasible, our design allowed for a comprehensive partitioning of phenotypic variation that we used to explicitly test for mitonuclear epistasis between two nuclear genomes and 22 mtDNAs. We modified the environment with rotenone, a natural inhibitor of mitochondrial Complex I– the largest OXPHOS complex containing seven mitochondrial and about 36 nuclear-encoded proteins [34]. This allowed us to challenge both the nuclear and mitochondrial genomes in our experiment and further partition phenotypic variation into environmental effects, GxE, and GxGxE interactions. The design was strengthened by duplicating the build of each mitonuclear combination and reproducibility was confirmed across experimental blocks. Significant main effects of the nuclear genome were detected across all four traits, while mtDNA and rotenone treatment effects were trait-dependent, confirming that variation in both genomes and environmental stressors jointly shaped phenotypic outcomes.

We found significant second order mitochondria-by-environment (GxE) interaction effects that influenced climbing velocity and development time. Indeed, for development time, the effect size of the mitochondrial GxE interaction was even greater than the environmental effect alone. It has been predicted that when exposed to novel and/or stressful environments (such as rotenone inhibiting OXPHOS through Complex I), GxE effects may be amplified because genotypes will respond unpredictably relative to their reactions to familiar environments [35]. The mitochondrial by rotenone interaction observed here may indicate cryptic mitochondrial variation, particularly for development time and male climbing velocity where interaction effects occur without mitochondrial main effects.

Across four traits measured in both sexes, we only found a single instance of a significant mitochondria-by-nuclear DNA interaction (i.e., mitonuclear epistasis or GxG)– male body weight. Body weight is strongly linked to metabolic function and the modulation of this trait by mitonuclear effects could be due to differences in OXPHOS efficiency. Beyond male weight, we found evidence of mitonuclear epistatic effects in male climbing at a less conservative significance threshold (*p* < 0.025), consistent with the high metabolic demands of this behavior [36]. Mitonuclear epistatic effects were also primarily driven by a few of the mtDNAs sampled, rather than a general property of the entire panel (Supplementary Figure 6). Previous studies have used covariation in evolutionary rates of mitochondrial and nuclear OXPHOS genes as strong evidence for mitonuclear coevolution in mammals [19], insects [37], and bivalves [18]. Therefore, disrupting this coevolved relationship may only be impactful for traits with high metabolic and OXPHOS demand or particular mitochondrial genotypes, reflecting the idiosyncratic nature of mitonuclear interactions found in both our study and others [38, 39].

Consistent with previous reports of context-dependent mitonuclear epistasis [40, 41, 42, 43, 44], our results indicate that GxG interactions vary across environments and sexes. In particular, we detected significant third-order interaction effects with the environment (GxGxE) and with sex (GxGxSex) in the climbing trait. Classical theory predicts that haploid, uniparentally inherited mtDNA is unlikely to maintain polymorphism unless fitness effects are finely balanced [45, 46]. However, context-dependent selection arising from sex, environment, or nuclear background may promote the maintenance of mtDNA variation through asymmetric fitness effects [47, 48, 49]. Our finding of both GxGxE and GxGxSex interactions may provide a mechanism for prolonging the polymorphic sojourn time of non-neutral mtDNA variants. Shifting selective advantages across contexts could buffer these variants against fixation compared to mtDNA evolving in a homogeneous environment.

Remarkably, across all significant mtDNA main effects, second- and third-order interactions, we found that the two outgroup species, *D. sim* and *D. yak*, consistently showed intermediate values relative to the two *D. mel* groups despite being paired with mismatched *D. mel* nuclear genomes. This result is surprising as it contrasts sharply with predictions from the nuclear compensation model, which suggests that nuclear alleles evolve to counteract deleterious mitochondrial mutations [11]—a form of coevolution expected to break down in interspecific mismatches. Furthermore, analyses of 53 nuclear-encoded OXPHOS genes in *Drosophila* showed that most are under purifying selection, with only three Complex I genes exhibiting signs of positive selection (Supplementary Table 15) [50, 51]. Our work bridges these molecular findings with phenotypic evidence, demonstrating that despite extensive amino acid divergence between species, mitonuclear communications have been maintained, limiting phenotypic differentiation across species boundaries. These results extend earlier work showing stronger within-species mitonuclear fitness effects [52], revealing that even across roughly 10 million years of divergence between *D. mel* and *D. yak*, nuclear compensation has not impacted these traits.

We propose that this observation of “excess polymorphism” relative to divergence is consistent with a model of stabilizing selection acting on mitonuclear communication, a pattern that is evident in other complex traits [7]. We found that the more divergent mtDNAs exhibited intermediate trait values as well as weaker interaction effects across environments for both GxE and GxGxE effects relative to the interspecific genotypes. These results are consistent with stabilizing selection for an optimal phenotype constraining divergence and promoting greater phenotypic stability across environmental gradients. To achieve this stability, purifying selection has likely prevented the fixation of harmful mutations between *D. mel, D. sim*, and *D. yak*. When we compared physical locations of amino acid changes in Complex I across *D. mel, D. sim* and *D. yak*, we found that within-species variants were enriched at protein-protein contact sites with the nuclear genome (Figure 5). Within-species variants also showed a higher proportion of radical AA changes (Supplementary Table 11), suggesting that most mtDNA variation within populations is mildly deleterious and transient and large-effect AA changes are purged before they can be fixed. This contrasts theory that suggests that purifying selection is weaker in the mitochondrial genome due to its reduced effective population size [53]. Our findings, together with evidence from *D. melanogaster* showing comparable mitochondrial and nuclear purifying selection [54] and consistently low mtDNA dN/dS values across the Genus *Drosophila* [55], support efficient purifying selection acting on mtDNA in this system.

Finally, our study addresses the Mother’s Curse hypothesis, which posits that maternal inheritance of mtDNA allows male-harming mutations to accumulate because selection acts only through female fitness [31, 32]. Cosmides & Tooby [56] presented early evidence for this selective asymmetry, showing that some mitochondrial diseases were more pronounced in males. By placing different mitochondrial haplotypes on a common nuclear background and measuring complex traits in both sexes, our design provides an effective empirical test of this hypothesis. Across the three traits we measured, we found no evidence for Mother’s Curse in either its strong or weak form, despite a study design that specifically addressed previous concerns about mtDNA sampling and reproducibility [33]. The absence of Mother’s Curse in our traits, combined with evidence of strong purifying or stabilizing selection on mtDNA, suggests that selection acting through females may indirectly constrain the accumulation of male-harming mutations.

As with any study of this scope, design constraints warrant consideration when interpreting our findings. First, our observation of large nuclear genome effects are limited to the two backgrounds that we sampled. Laboratory lines (here, DGRP375 and OreR) have been adapted to the laboratory environment for many generations and may respond to environmental perturbation differently than natural populations but were chosen as stocks that are fully sequenced and accessible to other researchers. We favored this decision rather than using an arbitrary line collected from the wild. Further, the ∼654,000 SNPs differing between these lines is a conservative estimate of nuclear divergence relative to the expected 1.8 million differences between lines collected from the wild (*π* ≈ 0.01). As such it seems likely that mitonuclear GxG and GxE could be more pronounced than what we have quantified here. We acknowledge that only a single mtDNA haplotype from each outgroup was used in this study; however, our aim was to assess whether increasingly divergent outgroups produced proportionally scaled effects of disrupted mitonuclear coadaptation and reasoned that one representative of each group was sufficient to capture the relevant amino acid divergence. Previous studies have used multiple mtDNAs from *D. simulans* in *D. melanogaster* backgrounds [52, 42], but no study has added *D. yakuba* to these kinds of analyses. Given that *D. yakuba* is ∼10 million years diverged from *D. melanogaster*, this provides a particularly robust test of the hypothesis that mitonuclear coadaptation should be disrupted by “foreign” mtDNAs [58, 11]. Environmental variation was captured along a single axis, a low dose of the insecticide rotenone. It may be the case that this is an unusually stressful environment, though we find that there is no impact on survival (Supplementary Tables 4–5) and where there are impacts on traits, the distributions overlap between the two diets (Figure 2), suggesting that this is instead a mild perturbation to the environment. Rotenone may not impact the full breath of possible mitonuclear interactions as it strictly inhibits Complex I; therefore, further work exploring how other environments interact with mitonuclear variation will be valuable for assessing how robust our findings are across contexts.

Collectively, our results demonstrate that mtDNA and its interactions with nuclear genetic variation and external environments are significant factors in the distribution of phenotypic variation in natural populations. Moreover, the very limited influence of divergent mtDNAs on these fitness related traits in *Drosophila* support a model of stabilizing selection acting on mitonuclear interactions that offer a distinct contrast to models that implicate mitonuclear interactions in adaptive evolution and speciation. The lack of major mitonuclear incompatibilities in *Drosophila* may reflect its relatively large effective populations size (*N*_*e*_) [57], which prevents the fixation of deleterious mtDNA mutations through efficient purifying selection. In contrast, smaller *N*_*e*_ systems such as *Tigriopus* copepods [58] or *Xiphophorus* swordtails [17] may allow for more harmful variants to accumulate, increasing the risk of incompatibilities. This “variable purging hypothesis” of mitonuclear incompatibilities is relevant to both evolutionary and medical applications. Our findings that mtDNA polymorphisms can be more impactful that divergent mtDNAs has implications for the choice of donor mtDNAs used in mitochondrial replacement therapy. If segregating mtDNA variants within various human population are more likely to harbor deleterious mutations than mtDNAs from more divergent populations, the latter may be more likely to have purged disease-causing mutations. Alternatively, widespread mtDNA haplogroups may have larger historical effective population sizes and may be less likely to carry such mutations. Future analyses should evaluate the important role of effective population size on the tempo and mode of mitonuclear coevolution.

## Methods

### Mitonuclear Lines

Polymorphism in mtDNA was sampled from 20 female lines of *D. melanogaster* (*D. mel)*, 10 from Beijing, China and 10 from Zimbabwe. Nineteen of these lines were originally characterized as the “Global Diversity Lines” (GDLs) [59], while the Zim53 line from Zimbabwe was originally characterized in Ballard 2000 [60]. Also included in the mtDNA panel were two genotypes from other species: *D. simulans* (*D. sim*) Hawaii (mitochondrial haplotype *siI* [61], mitonuclear construction described in [52]) and *D. yakuba* (*D. yak*) (mitonuclear construction described in [62]). Together, we used a total of 22 mitochondrial haplotypes incorporating intraspecific polymorphism and deep interspecific mitochondrial divergence from reproductively isolated outgroups.

To construct the mitonuclear lines, we first removed *Wolbachia* from each female line by treating them with tetracycline for two generations. PCR assays post treatment and again 2 years later confirmed the absence of *Wolbachia*. Following this, we created mitonuclear genotypes using 10 generations of recurrent male backcrossing from each of the nuclear lines to virgin females from the mitochondrial lines. We will describe the mitonuclear genotype of these lines using the notation *mtDNA;nucDNA*. We used two common nuclear backgrounds: *D. mel* Oregon R (*OreR*) and *D. mel* DGRP375 (*375*). For each mitonuclear combination, the construction was replicated by maintaining two independent lines (“builds”). Virgin females from each build were collected and independently crossed to males from either *OreR* or *375* nuclear backgrounds, providing an additional source of biological replication and controlling for any random effects of the backcrossing process. Also included in the panel were two additional “unaltered” lines for each nuclear background retaining their own native mitochondria (*OreR;OreR* and *375;375*). In total, 22 mitochondrial haplotypes were backcrossed in two independent builds to two nuclear backgrounds, plus the two genetically unaltered controls, resulting in 22×2×2 + 2 = 90 lines that were assayed. A complete list of all genotypes is provided in Supplementary Table 16.

### Fly rearing

Backcross lines were maintained on standard *Drosophila* rearing food. The food contains 1.8 g agar, 5 g SAF yeast, 10.4 g yellow cornmeal, 22 g sucrose, 0.9 tegosept in 4.5 ml 95% ethanol, cooked in 200 mL of deionized water and larger batches were scaled up with the same ratios. The rotenone diet was the standard control food containing 25 *µ*M of rotenone, a concentration selected to yield enough surviving adults with similar ages to controls. Prior to the phenotype assays, each line was expanded in multiple culture bottles before sorting 20 mated females from each bottle onto either control or rotenone food. The 20 females were allowed to lay eggs on either the control or rotenone food for 48 hours before being removed. The offspring of these individuals were then sorted by sex in 6 replicate vials of each sex and scored for each trait while being maintained on their respective diets.

### Phenotypic Assays

We measured body weight, climbing velocity, and flight performance in single-sex groups of adults. Prior to sorting adults by sex, we measured development time of mixed-sex groups. Development time was scored for three replicate vials following the 48-hour egg lay. Each day after the parental females were removed, we counted the number of pupae that have entered pupal stage P12 or above (identified by visible darkened wings [63]) for a total of 11 days post egg lay. We consider the average time since egg lay to be overall development time score for each vial. Following the development time assay, adult animals from the replicate vials were sorted by sex and flipped to new food of the same diet they were grown on. A total of six replicate vials of 20 males and six vials of 20 females were used for the remainder of the phenotype assays.

The climbing velocity or “negative geotaxis” assay was scored for all six vials in three technical replicate drops of the climbing rig to initiate climbing. The automated software FreeClimber was used to quantify the climbing performance [26]. FreeClimber quantifies the average velocity of animals within each vial by using a local linear regression on the average *y*-position vs. time and returns the steepest (i.e., fastest), linear two-second section of the curve as the final velocity. We consider the average of the three technical replicates to be the final score for each vial providing 3×6=18 estimates of climbing speed for each genotype, sex, and treatment combination.

To measure flight performance, we used three of the six sorted vials for each sex since this assay is sacrificial and the remaining vials needed to be saved for the body weight assay. The flight assay measures the response to an abrupt drop into a flight column where higher landing height is indicative of a stronger flight genotype. The details of the assay have been previously described in [25]. Due to the nature of the assay, all vials were scored simultaneously and we do not have access to individual trait scores for each vial. The trait value for each build of a genotype was then the average landing height for all flies across the three replicate vials.

Animal body weight was scored by combining the remaining surviving animals from three replicate vials of males and females and dividing the pool evenly between three 1.5 mL microcentrifuge tubes. Each microcentrifuge tube was pre-weighed using a scale with a precision of 0.001 mg (1 *µ*g) and then reweighed with a sample of approximately 20 animals. The exact number of animals per tube was recorded since it varied among samples. To get a final body weight, the initial weight of the tube was subtracted from the final weight of the tube with the flies and divided by the number of flies in each tube to estimate an average weight per fly. Since the animals from the remaining vials were pooled to maintain a more uniform distribution of animals per tube, we considered the average of the score for the replicate tubes to be the genotype score.

The concept of “build” introduced by the two independent backcrosses for each line provided an additional level of biological replication for each phenotype. That is, where we combine biological replicates to achieve an aggregate phenotype score in the flight and weight assays, there is still replication at the level of the build as the same genotype is tested multiple times.

### Experimental Blocks and Reproducibility of Phenotype

Due to the size of the mitonuclear panel and the number of the traits measured, the lines were tested across seven initial experimental blocks. The first three blocks were larger and contained eight intraspecific mitonuclear genotypes with both builds for a total of 16 lines tested. The remainder of the experimental blocks were scaled down to four intraspecific mitonuclear genotypes per block with both builds for a total of eight lines per block. Lines that were unsuccessful (i.e. low yield) in their original block were retested in a different block and removed from the original block. Lines in block 1 were remeasured for development time in block 8 due to an initial inconsistency with how this assay was implemented. Six “reference” genotypes were tested in every block and were used to adjust each trait measurement for block effects. This included one of the builds of the lines *siI;OreR, siI;375, yak;OreR, yak;375* and the two unaltered lines *OreR;OreR* and *375;375* (no associated build). For completeness, both builds of *siI;OreR, siI;375, yak;OreR, yak;375* were tested in the larger blocks 1-3 and in block 9.

To test for the reproducibility of the development time, the climbing, and the body weight phenotypes across experimental block environments, a subset of the lines tested in blocks 1 and 2 were retested in a final block (block 9). A total of 16 mitonuclear genotypes with both builds were retested in this final block for a total of 32 retested lines. We then compared the measures obtained from the second observation to those from the original testing block environment. The block assignments for each line are given in Table 16.

### ANOVA Model of Phenotypic Response

To test for the effects of rotenone treatment and genotype-phenotype associations for both the mitochondrial and nuclear genotypes, we used an ANOVA mixed model for each phenotype independently. To correct for variation between blocks, we subtracted the mean of the six “reference” genotypes (*OreR;OreR, 375;375*), and four “build A” lines of *yak;OreR, yak;375, siI;OreR, siI;375*) by block (*Bl*), treatment (*T*), and sex (*S*) from each raw data point and added the treatment and sex mean across all blocks 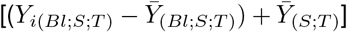. Including the interspecific (*D. yak* and *D. sim*) lines in the block correction helped prevent bias toward *D. mel* - specific responses, ensuring that block effects were removed based on the overall experimental panel rather than solely on *D. mel* performance.

The full model is given by:

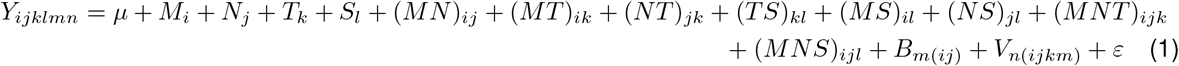

The model included the mitochondrial genotype (*M*), the nuclear genotype (*N*), treatment (*T*), and sex (*S*) as fixed effects as well as their interactions. Build (*B*), nested in the combination of the mitochondrial and nuclear genomes is included as a random effect. Build is a form of biological replication for each combination of mitochondrial and nuclear genotype. Within each build, there is a second level of replication that we model with random effects. For the climbing phenotypes there is an additional level of biological replication that comes from a shared environment of a vial (*V*) that is a random effect nested within build and treatment. Due to the way the treatment was applied (through diet), two treatments cannot be given to the same vial. Multiple measurements of each vial’s climbing velocity allows us to estimate the variance of this random effect. For the remainder of the phenotypes, we were only able to estimate the variance at the level of build (*B*). There is additionally an error term given by *ε*.

We also ran reduced models for each sex separately:

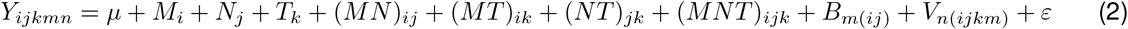

The development time phenotype does not have a sex term so only the reduced model was used. Additionally, since development time is affected by larval density, which was controlled for at the level of parental density rather than total egg count, there is an additional larval density term in the model which is approximated by the total pupal count in each vial divided by the total across all sets by treatment [*L* = total_*i*_/total_(*T*)_]. For the weight and flight phenotypes, averages at the level of build were used. We performed Type III ANOVA with Satterthwaite’s approximation for degrees of freedom. The type III ANOVA uses a partial sum of squares (SS) approach which adjusts the main effects to exclude the overlap with interaction terms. This means that the SS do not add to the total SS and can not be interpreted to partition the total variance of the model. To measure the relative importance of each fixed effect (effect size), we instead used a semi-partial *R*^2^ namely, 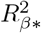, described in Jaeger et al., 2016 [64]. ANOVA models were implemented using the R v4.5.2 packages lme4 (v2.0-1, Supplementary References) and lmerTest (v3.2-1, Supplementary References); and 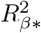 was calculated using the package r2glmm (v0.1.3, [64]). In Figure 2 we use the package emmeans (v2.0.2, Supplementary References) [3] to estimate the marginal means (or model-adjusted mean) for each factor of interest. We do this to obtain a more accurate portrayal of the effect being captured by the model.

To assess the stability of the interaction effects estimated by the ANOVA model, we refit the reduced models for each combination of trait/sex on subsets of mtDNA genotypes. We tested this for subsets of size *k* = (5, 10, 15, 20) where each level of *k* had 1000 unique (without replacement) randomly selected subsets of the 22 total mtDNAs. For *k* = 20, all 231 combinations were retested.

Note: When fitting the linear model for male weight, the random effect of build was estimated as having a near-zero variance, indicating that our data did not support among-build variation beyond residual error. This likely reflected limited replication and/or noisy measurements (males are ∼ 30% smaller than females).

### AMMI Model of Phenotypic Response

Where we identified significant third-order interactions using the ANOVA model, we used an Additive Main effects and Multiplicative Interaction model (AMMI) to further explore patterns in the data. AMMI is a frequently used technique for understanding complex genotype by environment interactions in agricultural yield trials and is particularly useful as a visualization technique [28, 65]. This method first applies analysis of variance to partition the variation in the phenotype of interest in to genotype effects (G), environmental effects (E), and genotype-environment (GE) interaction effects, and then it uses principal components analysis to decompose the GE effect. Since AMMI models explicitly decompose the interaction effect by first subtracting G and E additive effects, we can use the resultant decomposition to visualize the interactions present in the data. The AMMI*N* model equation is:

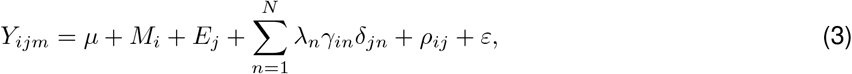

where each response *Y* is indexed by the mitochondrial genotype *i*, the environment *j*, and the build *m*. For these analyses, we used the adjusted *Y* values. Like in the ANOVA model, the mitochondrial main effect is given by *M*. Since the AMMI model is built to decompose GE interactions, we formed a single composite variable *E* for the “environment” that has factor levels for every combination of nuclear background and treatment (Ore:Control, Ore:Rotenone, 375:Control, and 375:Rotenone). This simplification is appropriate for visualizing patterns and aligns with prior AMMI applications incorporating additional environmental factors [28, 66]. The multiplicative part of the AMMI model uses PCA to decompose the remaining residual matrix after removing the *M* and *E* main effects into PCA axes 1 to *N*. The multiplicative parameters are: λ_*n*_ is the singular value for PCA axis *n, γ*_*in*_ is the mitochondrial genotype eigenvector for axis *n*, and *δ*_*jn*_ is the environment eigenvector for axis *n*. The AMMI multiplicative parameters λ^0.5^*γ*_*i*_ and λ^0.5^*δ*_*j*_ (for genotype and environment, respectively) are often referred to as “interaction PCA scores” (IPCA) to specify the distinction between typical PCA [28]. There are at most min(*M* − 1,*E* − 1) IPCA axes, but we are typically only interested in the first couple of axes (*n* < *N*). If not all axes are used, the remaining residual is included in *ρ*_*me*_. Additionally, we considered error at the replication level of build with the term *ε*_*ijm*_. Models were implemented using the R package agricolae (v1.3-7, Supplementary References).

### Fly Survival Assay

All lines were exposed to both standard control diets and diets containing 25 *µ*M of rotenone, a dose considerably lower than the lethal dose (750 *µ*M). However, to confirm that there was no significant effect of rotenone on the survival of the animals that may bias estimates of variance, we measured the egg to adult survival of 10 build A lines (Bei11, Bei54, *D. yak*, ZH42, and ZW133 mtDNA on OreR/DGRP nuclear backgrounds). Prior to the assay, each line was expanded in two culture bottles. Then, five females and five males were sorted onto eight replicate vials of each food type and laid eggs for ∼19 hours. Following the initial egg lay, animals were reciprocally flipped onto an additional eight replicate vials containing the other food type and again laid eggs for ∼19 hours. To assess total survival, we counted both the number of eggs laid in each vial and the total number of eclosed adults, sorted by sex. We used the following mixed linear model to test for the fixed effects of nuclear DNA (*N*), treatment (*T*), and egg count (*E*):

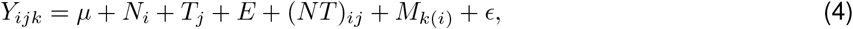

where *Y* is the total number of adults. Since we only tested one backcross build for each line, we cannot robustly measure the genetic effects of the mtDNA background. For this reason, we model the mtDNA effect (*M*) as a random effect nested within the nuclear background (similar to a line effect). We used the same model to assess the effect of sex, by fitting two additional models where *Y* is either the total count of males or females. We also tested for systematic differences in egg count due to a possible difference in preference for laying eggs on a control or treated food substrate using the following model:

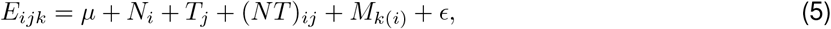

where the terms are the same as the survival model. It is an important distinction that in this model, treatment only refers to the environment where untreated females laid eggs and is not evaluating differences in fecundity due to animals grown in a rotenone environment. ANOVA models were implemented using lme4 (v2.0-1, Supplementary References) and lmerTest (v3.2-1, Supplementary References); 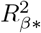 was calculated using the package r2glmm (v0.1.3, [64]); and we used emmeans package to estimate marginal means (v2.0.2, Supplementary References).

### Mother’s Curse tests

We consider two forms of this hypothesis. The weak form predicts that (1) males show greater variation in traits among the mitochondrial haplotypes than females and (2) there is no clear evidence of sexual antagonism (no linear relationship between male and female performance). The strong form predicts that (1) males and females both show variation across mitochondrial haplotypes and (2) the mutations between the haplotypes are sexually antagonistic in nature (negative linear relationship between female and male performance)[33].

To test predictions related to variation in trait performance, we average across technical replicates and biological replicates (build, vial) and calculate the variation due to mitochondrial genetic variation alone. We then compare the sex difference in this variation on the two nuclear backgrounds, OreR and DGRP375 and independently on both environments, control and rotenone. We sampled 1000 bootstraps (Supplemental Fig. 8) of the coefficient of variation (CV) for each sex and calculated the number of times the CV for males was less than or equal to the CV for females over the total number of bootstraps to obtain a *p*-value. Note that the Bei54 (375/Ore on rotenone) and ZW144 (Ore on rotenone) haplotypes were excluded from the analysis of the weight trait due to low sample sizes in males that artificially increased variance in males. To test predictions related to sexual antagonism between males and females, we calculated the correlation coefficient, *r*.

### mtDNA variants and phylogenetic signal

To call mtDNA variants, we retrieved DNA short reads from NCBI (PRJNA268111, [59] for GDL mtDNAs) and from lines sequenced by our research group (PRJNA1377152). Reads were aligned to the FlyBase *D*.*melanogaster* 6.32 reference using bwa (v0.7.17, Supplementary References). Variants were called using a standard GATK pipeline (excluding the AT-rich region of the mitochondrial genome), with base recalibration performed using known variants from the DGRP freeze 2 data [67], and final variant filtering applied according to GATK’s recommended hard-filtering parameters [6]. Variants were annotated with SNPeff (v4.3t, Supplementary References) to identify amino acid changes. The remaining mitochondrial genomes were downloaded from NCBI: Zim53 (KP843854.1, [68]), *D. yak* (NC_001322.1), and *D. sim* siI (AF200836.1). Changes to the amino acid sequence were obtained by aligning protein sequences with mafft (v7.505, Supplementary References) auto alignment and reporting changes in the sequence with custom python script. We retrieved the *D. mel* complex I protein structure from PDB (8B9Z, [29]) and used PyMOL (v3.1.6.1, Supplementary References) to identify protein-protein contact sites. Contact sites are defined as atoms from mtDNA-encoded proteins that are within 4Å of atoms from nucDNA-encoded proteins.

To evaluate the relationship between genetic relatedness and the trait measurements, we quantified phylogenetic signal. The major source of phylogenetic variation in our mitonuclear system was from the mitochondrial haplotypes, so we specifically quantified mitochondrial phylogenetic signal across an inferred mitochondrial phylogenetic tree. The filtered variants were used to create mitochondrial FASTA files for each sample using GATK’s ‘FastaAlternateReferenceMaker’. Multiple sequence alignment was performed using the mafft (v7.505, Supplementary References) ‘mafft-linsi’ algorithm and mitochondrial tree was inferred in RAxML-NG (v1.2.2, Supplementary References) using a majority-rule consensus tree derived from 1000 bootstrap replicates. We used the inferred phylogenetic tree to calculate Blomberg’s *K*, which estimates the strength of phylogenetic signal under a Brownian motion model of evolution [69]. To test if *K* is significantly different from zero, we sampled 1000 randomizations of the trait data across the phylogeny and calculated the number of times the randomized *K* is larger than the observed *K*. Dividing the proportion by 1000 yielded a *p*-value, which we then calculated for every combination of sex, treatment, nuclear background, and phenotype using the adjusted phenotype values. The R package phytools (v2.5-2, Supplementary References) was used to calculate both *K* and *p*-values.

## Data Availability

Code for data analyses are available at the GitHub site (https://github.com/leahdarwin/MitoGDL). Raw data files have been deposited to figshare. DNA short reads for a subset of the lines were deposited to the NCBI SRA under project number PRJNA1377152. The authors affirm that all analyzed data necessary for confirming the conclusions of the article are present within the article, figures, tables, supplementary information and GitHub site. Drosophila strains are available upon request from David Rand.

## Acknowledgments

The work was supported by grants from NIGMS: MIRA R35GM139607 to D. M. R., L. J. D. was supported by NIH T32 GM128596 and an NSF Graduate Fellowship (2023-2025), C. P. B. and O. C. M. were supported by NSF REU Division of Biological Infrastructure award 2150328, J. H. B. was supported by Brown University Undergraduate Teaching and Research Award. D. M. R. acknowledges support of COBRE award P20GM109035. We would also like to thank Chibuikem Nwizu and Ria Vinod for their helpful discussions regarding data analysis.

## Author Contributions

L. J. D. carried out the data analysis, wrote the manuscript, performed phenotype assays, and designed experiments. F. A. L. constructed the lines, performed phenotype assays and designed experiments. R. Z. B., J. H. B., C. P. B., A. D. G., J. D. L., and O. C. M. performed phenotype assays. Y. R. performed phenotype assays, edited the manuscript, designed experiments, and provided variant calling pipeline. D. M. R. conceptualized the study, performed phenotype assays, designed experiments, and edited the manuscript.

## References

[1] Visscher, P. M. et al. 10 years of GWAS discovery: biology, function, and translation. The American Journal of Human Genetics 101, 5–22 (2017).

[2] Hill, W. G., Goddard, M. E. & Visscher, P. M. Data and theory point to mainly additive genetic variance for complex traits. PLoS genetics 4, e1000008 (2008).

[3] Mackay, T. F. C. Epistasis and quantitative traits: using model organisms to study gene–gene interactions. Nature Reviews Genetics 15, 22–33 (2014).

[4] Zuk, O., Hechter, E., Sunyaev, S. R. & Lander, E. S. The mystery of missing heritability: genetic interactions create phantom heritability. Proceedings of the National Academy of Sciences 109, 1193–1198 (2012).

[5] Gimelfarb, A. Genotypic variation for a quantitative character maintained under stabilizing selection without mutations: epistasis. Genetics 123, 217–227 (1989).

[6] Barton, N. H. & Turelli, M. Effects of genetic drift on variance components under a general model of epistasis. Evolution; International Journal of Organic Evolution 58, 2111–2132 (2004).

[7] Mackay, T. F. C. Mutations and quantitative genetic variation: lessons from Drosophila. Philosophical Transactions of the Royal Society of London. Series B, Biological Sciences 365, 1229–1239 (2010).

[8] Sloan, D. B. et al. Cytonuclear integration and co-evolution. Nature Reviews Genetics 19, 635–648 (2018).

[9] Wolff, J. N., Ladoukakis, E. D., Enríquez, J. A. & Dowling, D. K. Mitonuclear interactions: evolutionary consequences over multiple biological scales. Philosophical Transactions of the Royal Society B: Biological Sciences 369, 20130443 (2014).

[10] Rand, D. M. & Mossman, J. A. Mitonuclear conflict and cooperation govern the integration of genotypes, phenotypes and environments. Philosophical Transactions of the Royal Society of London. Series B, Biological Sciences 375, 20190188 (2020).

[11] Rand, D. M., Haney, R. A. & Fry, A. J. Cytonuclear coevolution: the genomics of cooperation. Trends in Ecology & Evolution 19, 645–653 (2004).

[12] Hill, G. E. Mitonuclear compensatory coevolution. Trends in Genetics 36, 403–414 (2020).

[13] Pereira, R. J., Lima, T. G., Pierce-Ward, N. T., Chao, L. & Burton, R. S. Recovery from hybrid breakdown reveals a complex genetic architecture of mitonuclear incompatibilities. Molecular Ecology 30, 6403–6416 (2021).

[14] Nguyen, T. H. M. et al. Mapping mitonuclear epistasis using a novel recombinant yeast population. PLOS Genetics 19, e1010401 (2023).

[15] Bettinazzi, S. et al. Assessing the role of mitonuclear interactions on mitochondrial function and organismal fitness in natural Drosophila populations. Evolution Letters 8, 916–926 (2024).

[16] Chang, C.-C., Rodriguez, J. & Ross, J. Mitochondrial–nuclear epistasis impacts fitness and mitochondrial physiology of interpopulation Caenorhabditis briggsae hybrids. G3: Genes, Genomes, Genetics 6, 209–219 (2016).

[17] Robles, N. V. et al. Admixture mapping reveals evidence for multiple mitonuclear incompatibilities in swordtail fish hybrids Molecular Ecology 34, e70106 (2025).

[18] Piccinini, G. et al. Mitonuclear coevolution, but not nuclear compensation, drives evolution of OXPHOS complexes in bivalves. Molecular Biology and Evolution 38, 2597–2614 (2021).

[19] Weaver, R. J., Rabinowitz, S., Thueson, K. & Havird, J. C. Genomic signaturess of mitonuclear coevolution in mammals. Molecular Biology and Evolution 39, msac233 (2022).

[20] Zwonitzer, K. D. et al. Genome copy number predicts extreme evolutionary rate variation in plant mitochondrial DNA. Proceedings of the National Academy of Sciences 121, e2317240121 (2024).

[21] Sloan, D. B., Havird, J. C. & Sharbrough, J. The on-again, off-again relationship between mitochondrial genomes and species boundaries. Molecular Ecology 26, 2212–2236 (2017).

[22] Gupta, R. C., Milatovic, D. Biomarkers in Toxicology (Academic Press, Boston, 2014).

[23] Eyre-Walker, A. Mitochondrial replacement therapy: are mito-nuclear interactions likely to be a problem? Genetics 205, 1365–1372 (2017).

[24] McFarland, R. et al. Mitochondrial donation in a reproductive care pathway for mtDNA disease. New England Journal of Medicine 393, 461–468 (2025).

[25] Spierer, A. N. et al. Natural variation in the regulation of neurodevelopmental genes modifies flight performance in Drosophila. PLOS Genetics 17, e1008887 (2021).

[26] Spierer, A. N., Yoon, D., Zhu, C.-T. & Rand, D. M. FreeClimber: automated quantification of climbing performance in Drosophila. Journal of Experimental Biology 224, jeb229377 (2021).

[27] Varela, M. et al. Analysis of a three-way interaction including multi-attributes. Australian Journal of Agricultural Research 57, 1185–1193 (2006).

[28] Gauch (Jr.), H. G. Statistical Analysis of Regional Yield Trials: AMMI Analysis of Factorial Designs (Elsevier Science, Amsterdam, 1992).

[29] Agip, A.-N. A., Chung, I., Sanchez-Martinez, A., Whitworth, A. J. & Hirst, J. Cryo-EM structures of mitochondrial respiratory complex I from Drosophila melanogaster. eLife 12, e84424 (2023).

[30] Zhang, J. Rates of conservative and radical nonsynonymous nucleotide substitutions in mammalian nuclear genes. Journal of Molecular Evolution 50, 56–68 (2000).

[31] Frank, S. A. & Hurst, L. D. Mitochondria and male disease. Nature 383, 224 (1996).

[32] Gemmell, N. J., Metcalf, V. J. & Allendorf, F. W. Mother’s curse: the effect of mtDNA on individual fitness and population viability. Trends in Ecology & Evolution 19, 238–244 (2004).

[33] Dowling, D. K. & Adrian, R. E. Challenges and prospects for testing the mother’s curse hypothesis. Integrative and Comparative Biology 59, 875–889 (2019).

[34] Sharma, L. K., Lu, J. & Bai, Y. Mitochondrial respiratory complex I: structure, function and implication in human diseases. Current medicinal chemistry 16, 1266–1277 (2009).

[35] Saltz, J. B. et al. Why does the magnitude of genotype-by-environment interaction vary? Ecology and Evolution 8, 6342–6353 (2018).

[36] Sujkowski, A. et al. Mito-nuclear interactions modify Drosophila exercise performance. Mitochondrion 47, 188–205 (2019).

[37] Yan, Z., Ye, G. & Werren, J. H. Evolutionary rate correlation between mitochondrial-encoded and mitochondria-associated nuclear-encoded proteins in insects. Molecular Biology and Evolution 36, 1022–1036 (2019).

[38] Edmands, S. Mother’s curse effects on lifespan and aging. Frontiers in Aging 5 (2024).

[39] Garlovsky, M. D. et al. Testing for age- and sex-specific mitonuclear epistasis in Drosophila. Evolution 79, 1568–1582 (2025).

[40] Hoekstra, L. A., Siddiq, M. A. & Montooth, K. L. Pleiotropic effects of a mitochondrial–nuclear incompatibility depend upon the accelerating effect of temperature in Drosophila. Genetics 195, 1129–1139 (2013).

[41] Zhu, C.-T., Ingelmo, P. & Rand, D. M. G!G!E for lifespan in Drosophila: mitochondrial, nuclear, and dietary interactions that modify longevity. PLoS Genetics 10, e1004354 (2014).

[42] Mossman, J. A., Biancani, L. M., Zhu, C.-T. & Rand, D. M. Mitonuclear epistasis for development time and its modification by diet in Drosophila. Genetics 203, 463–484 (2016).

[43] Immonen, E., Berger, D., Sayadi, A., Liljestrand-Rönn, J. & Arnqvist, G. An experimental test of temperature-dependent selection on mitochondrial haplotypes in Callosobruchus maculatus seed beetles. Ecology And Evolution 10, 11387–11398 (2023).

[44] Erić, P., Veselinović, M., Patenković, A., Tanasković, M., Kenig, B., Erić, K., Inđić, B., Stanovčić, S. & Jelić, M. Mechanisms Maintaining Mitochondrial DNA Polymorphisms: The role of mito-nuclear interactions, sex-specific selection, and genotype-by-environment interactions in Drosophila subobscura. Insects 16, 415 (2025).

[45] Hedrick, P. W. Genetic polymorphism in heterogeneous environments: a decade later. Annual Review of Ecology, Evolution, and Systematics 17, 535–566 (1986).

[46] Clark, A. G. Natural selection with nuclear and cytoplasmic transmission. I. A deterministic model. Genetics 107, 679–701 (1984).

[47] Babcock, C. S. & Asmussen, M. A. Effects of differential selection in the sexes on cytonuclear dynamics: life stages with sex differences. Genetics 149, 2063–2077 (1998).

[48] Rand, D. M., Clark, A. G. & Kann, L. M. Sexually antagonistic cytonuclear fitness interactions in Drosophila melanogaster. Genetics 159, 173–187 (2001).

[49] Turelli, M. & Barton, N. H. Polygenic variation maintained by balancing selection: pleiotropy, sex-dependent allelic effects and G ! E interactions. Genetics 166, 1053–1079 (2004).

[50] Clark, A. G. et al. Evolution of genes and genomes on the Drosophila phylogeny. Nature 450, 203–218 (2007).

[51] Larracuente, A. M. et al. Evolution of protein-coding genes in Drosophila. Trends in genetics: TIG 24, 114–123 (2008).

[52] Montooth, K. L., Meiklejohn, C. D., Abt, D. N. & Rand, D. M. Mitochondrial-nuclear epistasis affects fitness within species but does not contribute to fixed incompatibilities between species of Drosophila. Evolution; International Journal of Organic Evolution 64, 3364–3379 (2010).

[53] Lynch, M. & Blanchard, J. L. Deleterious mutation accumulation in organelle genomes. Genetica 102-103, 29–39 (1998).

[54] Cooper, B. S., Burrus, C. R., Ji, C., Hahn, M. W. & Montooth, K. L. Similar efficacies of selection shape mitochondrial and nuclear genes in both Drosophila melanogaster and Homo sapiens. G3: Genes, Genomes, Genetics 5, 2165–2176 (2015).

[55] Montooth, K. L., Abt, D. N., Hofmann, J. W. & Rand, D. M. Comparative genomics of Drosophila mtDNA: novel features of conservation and change across functional domains and lineages. Journal of Molecular Evolution 69, 94–114 (2009).

[56] Cosmides, L. M. & Tooby, J. Cytoplasmic inheritance and intragenomic conflict. Journal of Theoretical Biology 89, 83–129 (1981).

[57] Sprengelmeyer, Q. D. et al. Recurrent collection of Drosophila melanogaster from wild African environments and genomic insights into species history. Molecular Biology and Evolution 37, 627–638 (2020).

[58] Burton, R. S. The role of mitonuclear incompatibilities in allopatric speciation. Cellular and Molecular Life Sciences 79, 103 (2022).

[59] Grenier, J. K. et al. Global Diversity Lines–A five-continent reference panel of sequenced Drosophila melanogaster strains. G3 Genes, Genomes, Genetics 5, 593–603 (2015).

[60] Ballard, J. W. O. Comparative Genomics of Mitochondrial DNA in Members of the Drosophila melanogaster Subgroup. Journal of Molecular Evolution 51, 48–63 (2000).

[61] Ballard, J. W. O. Comparative genomics of mitochondrial DNA in Drosophila simulans. Journal of Molecular Evolution 51, 64–75 (2000).

[62] Ma, L., Clark, A. G. & Keinan, A. Gene-based testing of interactions in association studies of quantitative traits. PLOS Genetics 9, e1003321 (2013).

[63] Chyb, S. & Gompel, N. Atlas of Drosophila Morphology: Wild-type and Classical Mutants (Academic Press, San Diego, 2013).

[64] Jaeger, B. C. E., Lloyd J.,, D, Kalyan, & Sen, P. K. An R2 statistic for fixed effects in the generalized linear mixed model. Journal of Applied Statistics 44, 1086–1105 (2017).

[65] Gauch, H. G. Statistical analysis of yield trials by AMMI and GGE. Crop Science 46, 1488–1500 (2006).

[66] Kona, P. et al. AMMI and GGE biplot analysis of genotype by environment interaction for yield and yield contributing traits in confectionery groundnut. Scientific Reports 14, 2943 (2024).

[67] Huang, W. et al. Natural variation in genome architecture among 205 Drosophila melanogaster Genetic Reference Panel lines. Genome Research 24, 1193–1208 (2009).

[68] Wolff, J. N. et al. Evolutionary implications of mitochondrial genetic variation: mitochondrial genetic effects on OXPHOS respiration and mitochondrial quantity change with age and sex in fruit flies. Journal of Evolutionary Biology 29, 736–747 (2016).

[69] Blomberg, S. P., Garland Jr., T. & Ives, A. R. Testing for phylogenetic signal in comparative data: behavioral traits are more labile. Evolution 57, 717–745 (2003).

[70] Early, A. M. & Clark, A. G. Monophyly of Wolbachia pipientis genomes within Drosophila melanogaster: geographic structuring, titre variation and host effects across five populations. Molecular Ecology 22, 5765–5778 (2013).

