## Supplemental Information for "Genetic and environmental interactions outweigh mitonuclear coevolution for complex traits in *Drosophila*"

#### Supplementary Information

#### Supplementary Figures

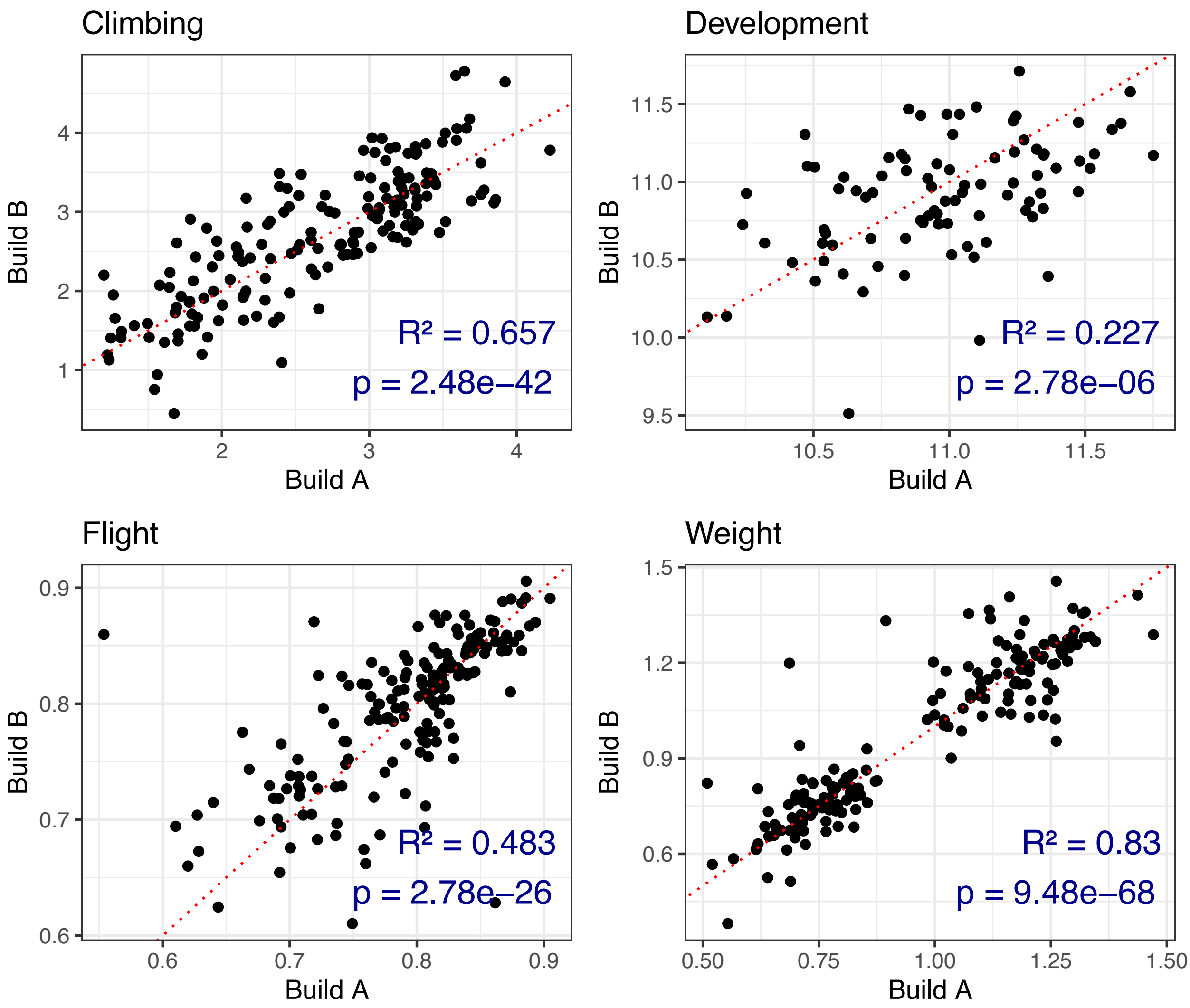


Supplementary Fig. 1: **Build-build correlations for each trait.** $R^2$ and $p$ values are given for Pearson's correlation test.

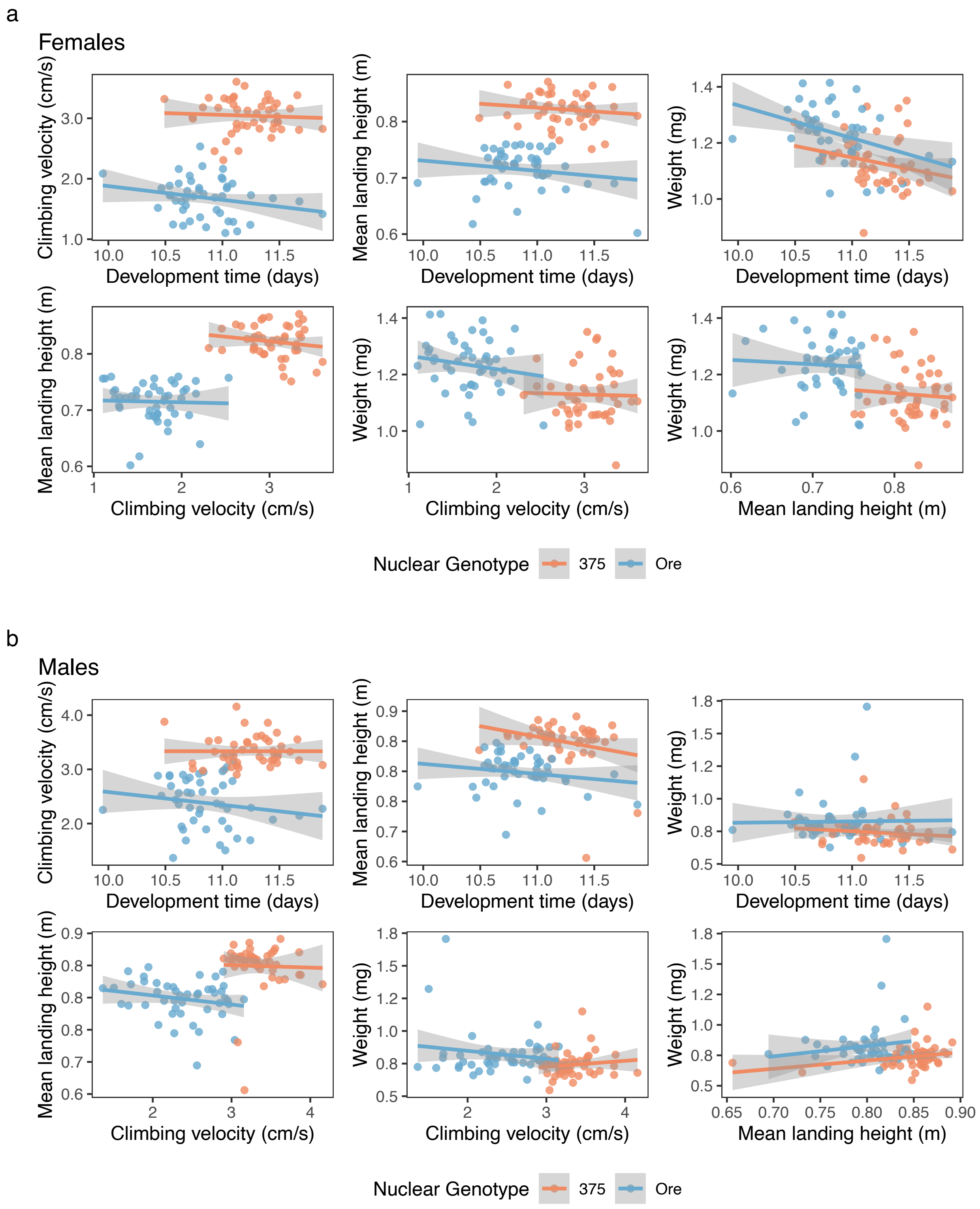


Supplementary Fig. 2: **Linear relationship between all phenotypes measured. a** Results for females. **b** Results for males. For each mitonuclear genotype and treatment, the values are averaged across all replicates and build. Due to large nuclear effect, the linear relationship is shown for each nuclear background independently.

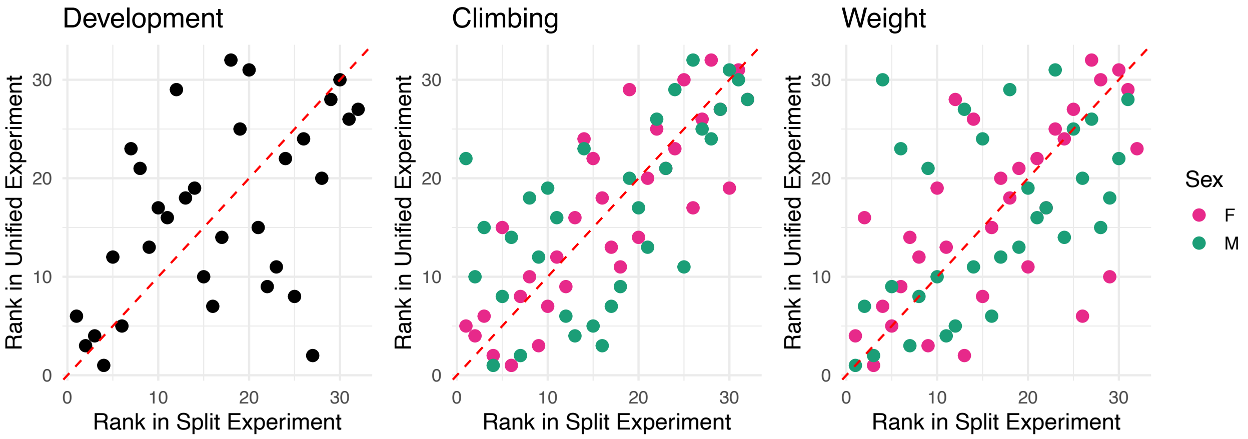


Supplementary Fig. 3: **Rank-rank plots for traits remeasured in set 9.** Rank-based correlations between trait measurements for lines measured within the same block or in different blocks. Males and females are plotted separately and development time was not scored by sex.

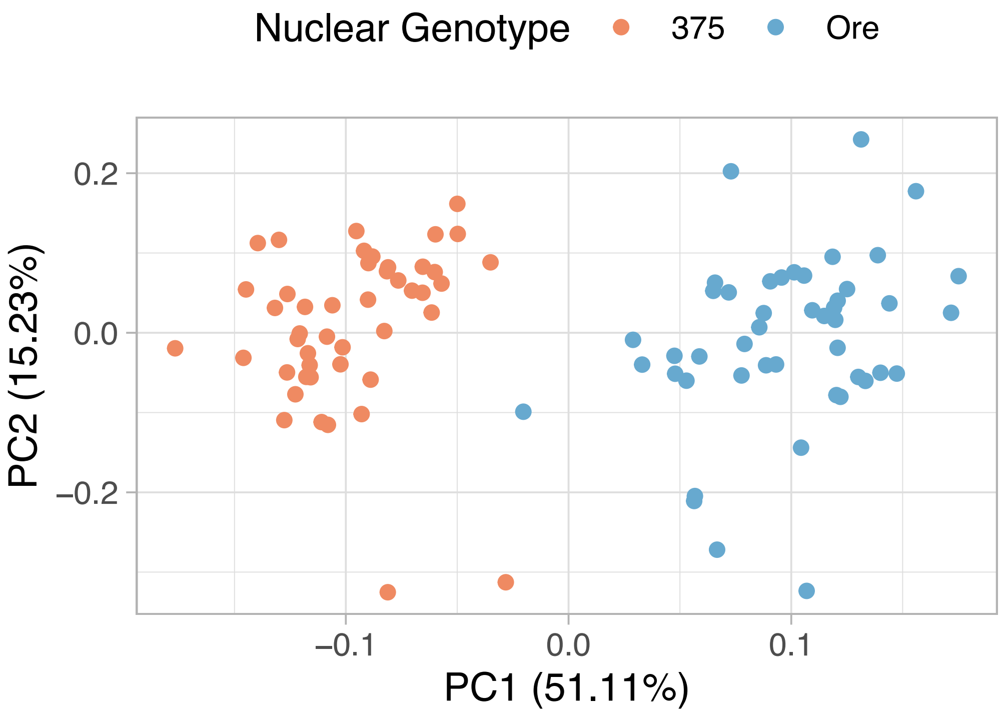


Supplementary Fig. 4: **PCA of phenotype space for males and females.** Values are averaged across builds and testing blocks if applicable and are unadjusted. The nuclear genotype explains 51.11% of the variance as seen along the first PC.

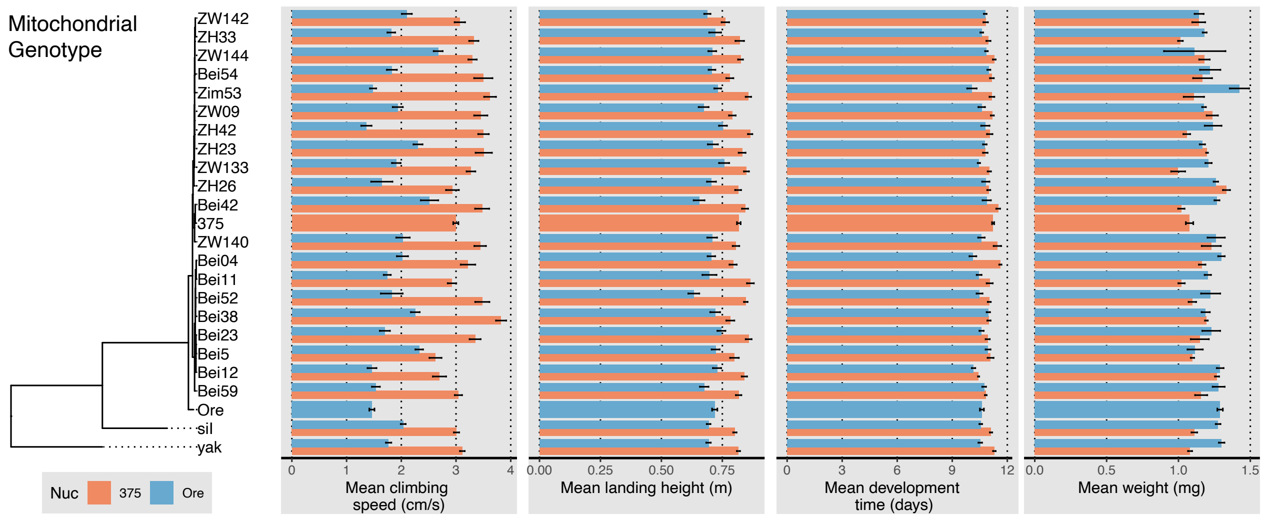


Supplementary Fig. 5: **Relationship between mitochondrial phylogeny and all phenotypes.** Mean trait values for both OreR and DGRP375 nuclear genotypes ordered by mitochondrial phylogenetic relatedness. Only females on control food are shown.

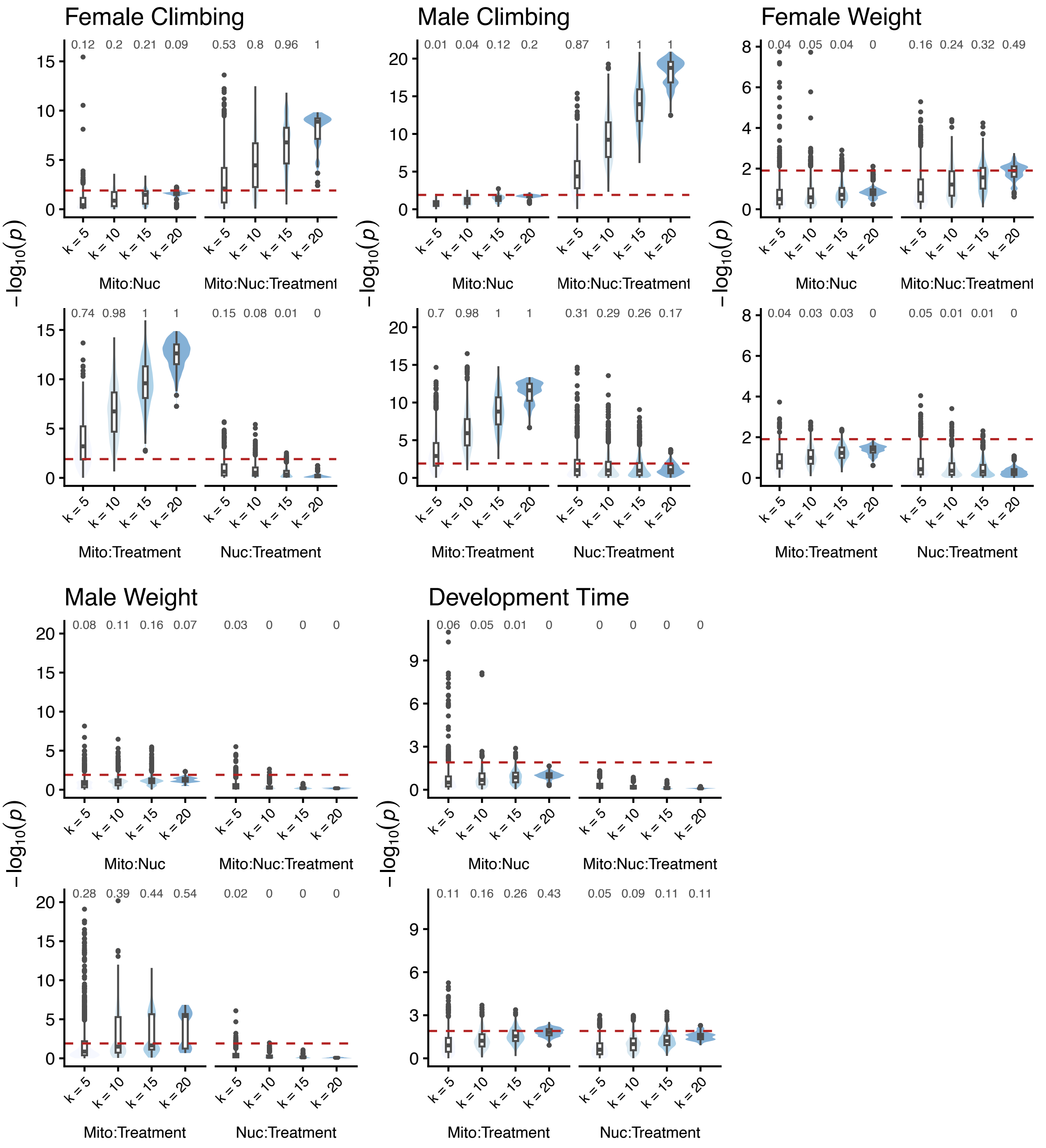


Supplementary Fig. 6: **Interaction effect stability across subsamples of $k$ mtDNAs.** For each sex/trait combination, the model is refit for 1000 resamples without replacement of the data for subsets of size $k$ mtDNAs of the 22 total mtDNAs ($k = 20$, enumerated all 232 combinations). The per-term distributions of the resultant p-values are given for each $k$ value tested. The red dotted line shows the significance threshold used for the original ANOVA tests and the number labels give the proportion of subsets where the threshold is passed.

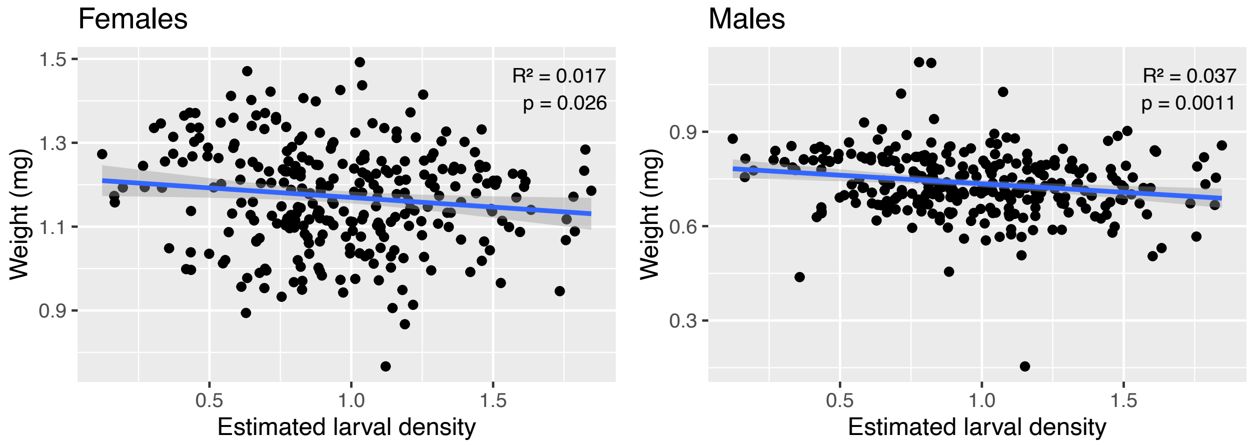


Supplementary Fig. 7: **Relationship between estimated larval density and weight measurement for females and males.** Where an estimate of larval density was made for each genotype/treatment/experimental block combination, we show the relationship between weight for females (left) and males (right). $R^2$ and $p$ values are given for Pearson's correlation test for each sex.

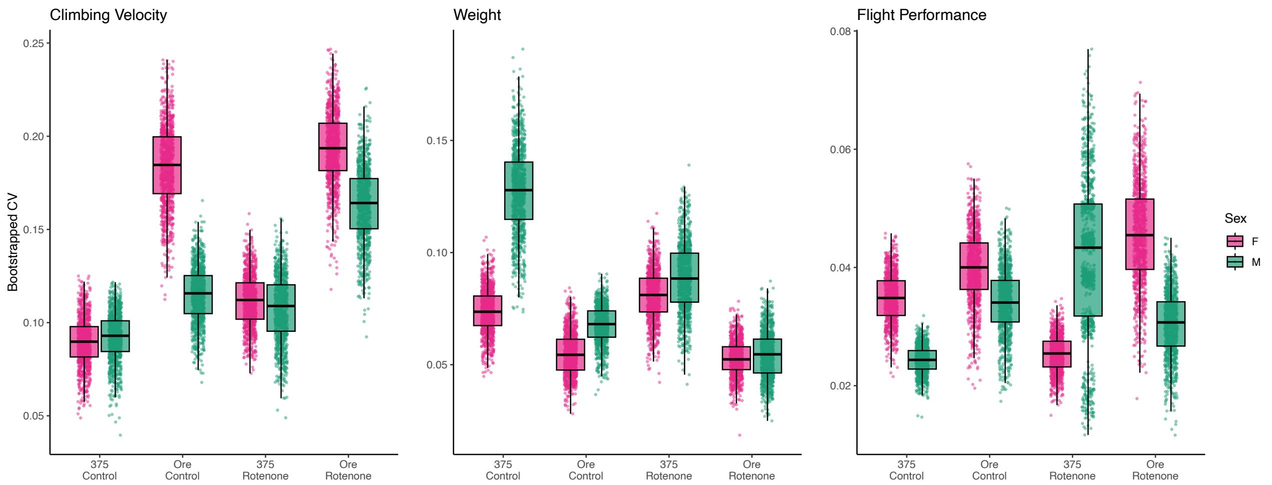


Supplementary Fig. 8: **Mother's Curse comparison of CV between sexes.** Bootstrapped (n=1000) distributions of the coefficient of variation (CV) for each trait. Comparisons between females (F) and males (M) are given for each combination of nuclear background (OreR, DGRP375) and treatment environment (control, rotenone).

### Supplementary Tables

Supplementary Table 1: **Estimation of the trait-trait Pearson correlation coefficient $r$ in females.** For each combinations of traits, the correlation coefficient $r$ and 95% confidence interval ($r_{lower}$,$r_{upper}$) is given. Trait values were unadjusted and tests were performed for each pair of phenotypes measured. Data is additionally stratified on nuclear genotype as shown in the Nuc column. The Bonferroni corrected significance threshold is $\alpha = \frac{0.05}{\text{12 comparisons}} = 0.004$ for the given $p$-value testing the alternative hypothesis that $r$ is different from 0.

| Phenotype 1 | Phenotype 2 | Nuc | $r$ | $r_{lower}$ | $r_{upper}$ | $p$ |
|---|---|---|---|---|---|---|
| Development time (days) | Climbing velocity (cm/s) | 375 | -0.056 | -0.341 | 0.238 | 0.713 |
| Development time (days) | Climbing velocity (cm/s) | Ore | -0.231 | -0.488 | 0.064 | 0.123 |
| Development time (days) | Mean landing height (m) | 375 | -0.133 | -0.407 | 0.164 | 0.379 |
| Development time (days) | Mean landing height (m) | Ore | -0.166 | -0.436 | 0.130 | 0.269 |
| Development time (days) | Weight (mg) | 375 | -0.242 | -0.497 | 0.052 | 0.106 |
| Development time (days) | Weight (mg) | Ore | -0.400 | -0.619 | -0.125 | 0.006 |
| Climbing velocity (cm/s) | Mean landing height (m) | 375 | -0.164 | -0.433 | 0.133 | 0.277 |
| Climbing velocity (cm/s) | Mean landing height (m) | Ore | -0.032 | -0.319 | 0.261 | 0.833 |
| Climbing velocity (cm/s) | Weight (mg) | 375 | -0.027 | -0.315 | 0.266 | 0.860 |
| Climbing velocity (cm/s) | Weight (mg) | Ore | -0.160 | -0.431 | 0.136 | 0.287 |
| Mean landing height (m) | Weight (mg) | 375 | -0.069 | -0.352 | 0.226 | 0.650 |
| Mean landing height (m) | Weight (mg) | Ore | -0.058 | -0.343 | 0.236 | 0.701 |

Supplementary Table 2: **Estimation of the trait-trait Pearson correlation coefficient $r$ in males.** For each combinations of traits, the correlation coefficient $r$ and 95% confidence interval ($r_{lower}$,$r_{upper}$) is given. Trait values were unadjusted and tests were performed for each pair of phenotypes measured. Data is additionally stratified on nuclear genotype as shown in the Nuc column. The Bonferroni corrected significance threshold is $\alpha = \frac{0.05}{\text{12 comparisons}} = 0.004$ for the given $p$-value testing the alternative hypothesis that $r$ is different from 0.

| Phenotype 1 | Phenotype 2 | Nuc | $r$ | $r_{lower}$ | $r_{upper}$ | $p$ |
|---|---|---|---|---|---|---|
| Development time (days) | Climbing velocity (cm/s) | 375 | 0.000 | -0.290 | 0.291 | 0.998 |
| Development time (days) | Climbing velocity (cm/s) | Ore | -0.171 | -0.439 | 0.126 | 0.257 |
| Development time (days) | Mean landing height (m) | 375 | -0.253 | -0.506 | 0.041 | 0.090 |
| Development time (days) | Mean landing height (m) | Ore | -0.183 | -0.449 | 0.114 | 0.225 |
| Development time (days) | Weight (mg) | 375 | -0.128 | -0.403 | 0.169 | 0.398 |
| Development time (days) | Weight (mg) | Ore | 0.019 | -0.273 | 0.307 | 0.901 |
| Climbing velocity (cm/s) | Mean landing height (m) | 375 | -0.028 | -0.316 | 0.265 | 0.854 |
| Climbing velocity (cm/s) | Mean landing height (m) | Ore | -0.213 | -0.474 | 0.082 | 0.155 |
| Climbing velocity (cm/s) | Weight (mg) | 375 | 0.132 | -0.165 | 0.406 | 0.383 |
| Climbing velocity (cm/s) | Weight (mg) | Ore | -0.162 | -0.432 | 0.135 | 0.282 |
| Mean landing height (m) | Weight (mg) | 375 | 0.268 | -0.024 | 0.518 | 0.072 |
| Mean landing height (m) | Weight (mg) | Ore | 0.148 | -0.148 | 0.421 | 0.325 |

Supplementary Table 3: **Spearman's rank correlation $\rho$ for retested lines.** Correlation coefficient $\rho$ and the 95% confidence interval ($\rho_{lower}$,$\rho_{upper}$) is given for traits for mitonuclear lines measured in different blocks and then again in the same block. The $p$-value given for the alternative hypothesis that $\rho$ is different from 0.

| Trait | Sex | $\rho$ | $\rho_{lower}$ | $\rho_{upper}$ | $p$ |
|---|---|---|---|---|---|
| Climbing | F | 0.842 | 0.699 | 0.921 | $<$0.001 |
| Climbing | M | 0.631 | 0.362 | 0.803 | $<$0.001 |
| Weight | F | 0.636 | 0.369 | 0.806 | $<$0.001 |
| Weight | M | 0.506 | 0.185 | 0.730 | 0.004 |
| Development | MF | 0.488 | 0.168 | 0.715 | 0.005 |

Supplementary Table 4: **ANOVA for survival on rotenone diet.** Analysis of variance for survival assay on a subset of 10 lines (see Methods). Three different models were tested, total count (males+females) and counts for males and females separately. For each term, the $F$-statistic, $p$-value, and semi-partial $R^2$ for fixed effects is given.

| Term | SS | MS | $df$ | $F$ | $p$ | $R^2_{\beta*}$ |
|---|---|---|---|---|---|---|
| **Total** | | | | | | |
| Nuc | 2306.7900 | 2306.7900 | 1 | 15.5626 | 0.00146 | 0.1602 |
| Treatment | 43.2036 | 43.2036 | 1 | 0.2915 | 0.59009 | 0.0017 |
| Egg Count | 59996.6128 | 59996.6128 | 1 | 404.7639 | < 0.001 | 0.7533 |
| Nuc:Treatment | 476.9487 | 476.9487 | 1 | 3.2177 | 0.07490 | 0.0186 |
| **Female** | | | | | | |
| Nuc | 1078.6226 | 1078.6226 | 1 | 18.4451 | <0.001 | 0.1274 |
| Treatment | 53.1665 | 53.1665 | 1 | 0.9092 | 0.342 | 0.0056 |
| Egg Count | 17894.3839 | 17894.3839 | 1 | 306.0040 | <0.001 | 0.6775 |
| Nuc:Treatment | 103.2756 | 103.2756 | 1 | 1.7661 | 0.186 | 0.0107 |
| **Male** | | | | | | |
| Nuc | 367.1076 | 367.1076 | 1 | 7.6412 | 0.0166 | 0.1024 |
| Treatment | 0.3123 | 0.3123 | 1 | 0.0065 | 0.9359 | 0.0000 |
| Egg Count | 14228.0723 | 14228.0723 | 1 | 296.1502 | <0.001 | 0.6952 |
| Nuc:Treatment | 138.0217 | 138.0217 | 1 | 2.8729 | 0.0922 | 0.0164 |

Supplementary Table 5: **Estimated marginal mean survivors.** Estimated marginal means (EMM) of total survivors from survival assay on a subset of 10 lines (see Methods) on control and rotenone diets. Egg Count is the grand mean across all factors. Means are given for models using the total count (male+female), female count, and male count.

| Treatment | Egg Count | EMM | SE | $df$ | Lower CI | Upper CI |
|---|---|---|---|---|---|---|
| **Total** | | | | | | |
| Control | 82.3375 | 55.5456 | 1.8843 | 13.9479 | 51.5027 | 59.5885 |
| Rotenone | 82.3375 | 54.4970 | 1.8968 | 14.3100 | 50.4369 | 58.5570 |
| **Female** | | | | | | |
| Control | | 28.5774 | 0.9446 | 20.7793 | 26.6116 | 30.5431 |
| Rotenone | | 27.4157 | 0.9544 | 21.5942 | 25.4342 | 29.3972 |
| **Male** | | | | | | |
| Control | | 26.9801 | 1.1950 | 12.3670 | 24.3850 | 29.5752 |
| Rotenone | | 27.0693 | 1.2014 | 12.6279 | 24.4660 | 29.6725 |

Supplementary Table 6: **ANOVA for egg count of control adults.** Analysis of variance for egg counts measured in the survival assay on a subset of 10 lines (see Methods). Parents were all grown on control diets, therefore the treatment effect measures preference for egg lay on food type. For each term, the $F$-statistic, $p$-value, and semi-partial $R^2$ for fixed effects is given.

| Term | SS | MS | $df$ | $F$ | $p$ | $R^2_{\beta*}$ |
|---|---|---|---|---|---|---|
| **Egg Count** | | | | | | |
| Nuc | 23300.2176 | 23300.2176 | 1 | 22.3853 | 0.00148 | 0.5364 |
| Treatment | 3090.8397 | 3090.8397 | 1 | 2.9695 | 0.08694 | 0.0160 |
| Nuc:Treatment | 17.0479 | 17.0479 | 1 | 0.0164 | 0.89834 | 0.0001 |

Supplementary Table 7: **ANOVA for traits in females.** Analysis of variance for sex-stratified model of female animals for the climbing, flight, and weight traits. The Bonferroni corrected significance threshold is $\alpha = \frac{0.05}{\text{4 traits}} = 0.025$. The given $R^2$ value is the semi-partial $R^2$ for fixed effects.

| Trait | Term | SS | MS | *df* | $F$ | $p$ | $R^2_{\beta*}$ |
|---|---|---|---|---|---|---|---|
| Climbing | Mito | 25.3589 | 1.2076 | 21 | 2.4999 | 0.0053 | 0.0500 |
| Climbing | Nuc | 305.8063 | 305.8063 | 1 | 633.0724 | < 0.001 | 0.3740 |
| Climbing | Treatment | 74.3476 | 74.3476 | 1 | 153.9126 | < 0.001 | 0.0334 |
| Climbing | Mito:Nuc | 20.2900 | 0.9662 | 21 | 2.0002 | 0.0267 | 0.0404 |
| Climbing | Mito:Treatment | 57.1195 | 2.7200 | 21 | 5.6308 | < 0.001 | 0.0276 |
| Climbing | Nuc:Treatment | 0.0001 | 0.0001 | 1 | 0.0003 | 0.9872 | 0.0000 |
| Climbing | Mito:Nuc:Treatment | 42.3851 | 2.0183 | 21 | 4.1783 | < 0.001 | 0.0215 |
| Flight | Mito | 0.0248 | 0.0012 | 21 | 1.7847 | 0.0525 | 0.2464 |
| Flight | Nuc | 0.2414 | 0.2414 | 1 | 364.3893 | <0.001 | 0.7606 |
| Flight | Treatment | 0.0000 | 0.0000 | 1 | 0.0552 | 0.8153 | 0.0002 |
| Flight | Mito:Nuc | 0.0190 | 0.0009 | 21 | 1.3641 | 0.1895 | 0.2003 |
| Flight | Mito:Treatment | 0.0307 | 0.0015 | 21 | 2.2032 | 0.0139 | 0.1676 |
| Flight | Nuc:Treatment | 0.0028 | 0.0028 | 1 | 4.2664 | 0.0448 | 0.0183 |
| Flight | Mito:Nuc:Treatment | 0.0231 | 0.0011 | 21 | 1.6578 | 0.0792 | 0.1312 |
| Weight | Mito | 0.9568 | 0.0456 | 21 | 4.1333 | <0.001 | 0.2378 |
| Weight | Nuc | 0.6279 | 0.6279 | 1 | 56.9588 | <0.001 | 0.1577 |
| Weight | Treatment | 0.0003 | 0.0003 | 1 | 0.0311 | 0.8601 | 0.0000 |
| Weight | Mito:Nuc | 0.3498 | 0.0167 | 21 | 1.5113 | 0.1346 | 0.1218 |
| Weight | Mito:Treatment | 0.3802 | 0.0181 | 21 | 1.6423 | 0.0354 | 0.0487 |
| Weight | Nuc:Treatment | 0.0044 | 0.0044 | 1 | 0.3983 | 0.5282 | 0.0006 |
| Weight | Mito:Nuc:Treatment | 0.4293 | 0.0204 | 21 | 1.8547 | 0.0116 | 0.0546 |

Supplementary Table 8: **ANOVA for traits in males.** Analysis of variance for sex-stratified model of male animals for the climbing, flight, and weight traits. The Bonferroni corrected significance threshold is $\alpha = \frac{0.05}{\text{4 traits}} = 0.025$. The given $R^2$ value is the semi-partial $R^2$ for fixed effects.

| Trait | Term | SS | MS | *df* | $F$ | $p$ | $R^2_{\beta*}$ |
|---|---|---|---|---|---|---|---|
| Climbing | Mito | 11.1825 | 0.5325 | 21 | 0.8205 | 0.6815 | 0.0206 |
| Climbing | Nuc | 121.0072 | 121.0072 | 1 | 186.4606 | <0.001 | 0.1827 |
| Climbing | Treatment | 80.1409 | 80.1409 | 1 | 123.4896 | <0.001 | 0.0238 |
| Climbing | Mito:Nuc | 29.6560 | 1.4122 | 21 | 2.1761 | 0.0150 | 0.0488 |
| Climbing | Mito:Treatment | 71.5921 | 3.4091 | 21 | 5.2532 | <0.001 | 0.0242 |
| Climbing | Nuc:Treatment | 2.1675 | 2.1675 | 1 | 3.3399 | 0.0679 | 0.0007 |
| Climbing | Mito:Nuc:Treatment | 102.6115 | 4.8863 | 21 | 7.5293 | <0.001 | 0.0328 |
| Flight | Mito | 0.0308 | 0.0015 | 21 | 1.0869 | 0.395 | 0.1348 |
| Flight | Nuc | 0.1007 | 0.1007 | 1 | 74.5448 | <0.001 | 0.3373 |
| Flight | Treatment | 0.0248 | 0.0248 | 1 | 18.3899 | <0.001 | 0.0928 |
| Flight | Mito:Nuc | 0.0389 | 0.0019 | 21 | 1.3721 | 0.185 | 0.1645 |
| Flight | Mito:Treatment | 0.0388 | 0.0018 | 21 | 1.3678 | 0.188 | 0.1376 |
| Flight | Nuc:Treatment | 0.0008 | 0.0008 | 1 | 0.6256 | 0.433 | 0.0035 |
| Flight | Mito:Nuc:Treatment | 0.0171 | 0.0008 | 21 | 0.6031 | 0.894 | 0.0657 |
| Weight | Mito | 1.5973 | 0.0761 | 21 | 6.8384 | <0.001 | 0.1991 |
| Weight | Nuc | 0.2251 | 0.2251 | 1 | 20.2407 | <0.001 | 0.0338 |
| Weight | Treatment | 0.0049 | 0.0049 | 1 | 0.4400 | 0.507 | 0.0008 |
| Weight | Mito:Nuc | 0.6253 | 0.0298 | 21 | 2.6772 | <0.001 | 0.0887 |
| Weight | Mito:Treatment | 0.8410 | 0.0400 | 21 | 3.6005 | <0.001 | 0.1157 |
| Weight | Nuc:Treatment | 0.0006 | 0.0006 | 1 | 0.0506 | 0.822 | 0.0001 |
| Weight | Mito:Nuc:Treatment | 0.1883 | 0.0090 | 21 | 0.8063 | 0.714 | 0.0285 |

Supplementary Table 9: **ANOVA for full model of all traits.** Analysis of variance for the model with sex as a term for the climbing, flight, weight, and development time traits. The corrected significance threshold is $\alpha = \frac{0.05}{4 \text{ traits}} = 0.025$. The development time trait was not scored for each sex and does not have a sex term. Additionally, development time has a larval density term associated with how the assay was conducted. The given $R^2$ value is the semi-partial $R^2$ for fixed effects.

| Trait | Term | SS | MS | $df$ | $F$ | $p$ | $R^2_{\beta*}$ |
|---|---|---|---|---|---|---|---|
| Climb | Mito | 15.1929 | 0.7235 | 21 | 1.2082 | 0.2916 | 0.0222 |
| Climb | Nuc | 284.6815 | 284.6815 | 1 | 475.4245 | <0.001 | 0.2710 |
| Climb | Treatment | 154.9403 | 154.9403 | 1 | 258.7538 | <0.001 | 0.0282 |
| Climb | Sex | 487.4681 | 487.4681 | 1 | 814.0828 | <0.001 | 0.0622 |
| Climb | Mito:Nuc | 27.0453 | 1.2879 | 21 | 2.1508 | 0.0164 | 0.0319 |
| Climb | Mito:Treatment | 104.2621 | 4.9649 | 21 | 8.2914 | <0.001 | 0.0217 |
| Climb | Nuc:Treatment | 1.1459 | 1.1459 | 1 | 1.9136 | 0.1668 | 0.0002 |
| Climb | Treatment:Sex | 0.0559 | 0.0559 | 1 | 0.0933 | 0.7600 | 0.0000 |
| Climb | Mito:Sex | 87.7973 | 4.1808 | 21 | 6.9821 | <0.001 | 0.0118 |
| Climb | Nuc:Sex | 83.5805 | 83.5805 | 1 | 139.5813 | <0.001 | 0.0113 |
| Climb | Mito:Nuc:Treatment | 87.5380 | 4.1685 | 21 | 6.9614 | <0.001 | 0.0185 |
| Climb | Mito:Nuc:Sex | 108.7268 | 5.1775 | 21 | 8.6465 | <0.001 | 0.0146 |
| Flight | Mito | 0.0316 | 0.0015 | 21 | 1.4736 | 0.1374 | 0.1372 |
| Flight | Nuc | 0.2952 | 0.2952 | 1 | 288.6215 | < 0.001 | 0.5984 |
| Flight | Treatment | 0.0134 | 0.0134 | 1 | 13.1249 | < 0.001 | 0.0351 |
| Flight | Sex | 0.2813 | 0.2813 | 1 | 275.0916 | < 0.001 | 0.4316 |
| Flight | Mito:Nuc | 0.0236 | 0.0011 | 21 | 1.0977 | 0.3848 | 0.1066 |
| Flight | Mito:Treatment | 0.0486 | 0.0023 | 21 | 2.2610 | 0.0022 | 0.1159 |
| Flight | Nuc:Treatment | 0.0033 | 0.0033 | 1 | 3.2638 | 0.0725 | 0.0090 |
| Flight | Treatment:Sex | 0.0116 | 0.0116 | 1 | 11.3713 | < 0.001 | 0.0304 |
| Flight | Mito:Sex | 0.0287 | 0.0014 | 21 | 1.3349 | 0.1588 | 0.0718 |
| Flight | Nuc:Sex | 0.0616 | 0.0616 | 1 | 60.2533 | < 0.001 | 0.1426 |
| Flight | Mito:Nuc:Treatment | 0.0190 | 0.0009 | 21 | 0.8864 | 0.6087 | 0.0489 |
| Flight | Mito:Nuc:Sex | 0.0424 | 0.0020 | 21 | 1.9755 | 0.0093 | 0.1027 |
| Weight | Mito | 0.8710 | 0.0415 | 21 | 3.6753 | <0.001 | 0.2295 |
| Weight | Nuc | 0.4218 | 0.4218 | 1 | 37.3799 | <0.001 | 0.1083 |
| Weight | Treatment | 0.0270 | 0.0270 | 1 | 2.3890 | 0.1224 | 0.0025 |
| Weight | Sex | 53.3014 | 53.3014 | 1 | 4723.3066 | <0.001 | 0.8318 |
| Weight | Mito:Nuc | 0.3864 | 0.0184 | 21 | 1.6305 | 0.0976 | 0.1181 |
| Weight | Mito:Treatment | 0.3358 | 0.0160 | 21 | 1.4168 | 0.0995 | 0.0304 |
| Weight | Nuc:Treatment | 0.0031 | 0.0031 | 1 | 0.2775 | 0.5984 | 0.0003 |
| Weight | Treatment:Sex | 0.1779 | 0.1779 | 1 | 15.7643 | <0.001 | 0.0162 |
| Weight | Mito:Sex | 0.5237 | 0.0249 | 21 | 2.2101 | 0.0013 | 0.0463 |
| Weight | Nuc:Sex | 0.1976 | 0.1976 | 1 | 17.5061 | <0.001 | 0.0180 |
| Weight | Mito:Nuc:Treatment | 0.2777 | 0.0132 | 21 | 1.1719 | 0.2667 | 0.0251 |
| Weight | Mito:Nuc:Sex | 0.3217 | 0.0153 | 21 | 1.3575 | 0.1289 | 0.0290 |
| Development | Mito | 5.3462 | 0.2546 | 21 | 1.7637 | 0.0630 | 0.1139 |
| Development | Nuc | 10.9876 | 10.9876 | 1 | 76.1200 | <0.001 | 0.2034 |
| Development | Treatment | 5.1530 | 5.1530 | 1 | 35.6987 | <0.001 | 0.0431 |
| Development | Larval density | 1.7335 | 1.7335 | 1 | 12.0095 | <0.001 | 0.0164 |
| Development | Mito:Nuc | 4.9106 | 0.2338 | 21 | 1.6200 | 0.0967 | 0.1148 |
| Development | Mito:Treatment | 5.5793 | 0.2657 | 21 | 1.8406 | 0.0123 | 0.0465 |
| Development | Nuc:Treatment | 0.7532 | 0.7532 | 1 | 5.2183 | 0.0226 | 0.0065 |
| Development | Mito:Nuc:Treatment | 2.1318 | 0.1015 | 21 | 0.7033 | 0.8325 | 0.0183 |

Supplementary Table 10: **Estimation of mtDNA phylogenetic signal.** Blomberg's $K$ values for mitochondrial signal across all sex, nuclear genotype, and treatment combinations. The given $p$-value is proportion of times a random permutation of the trait data resulted in a greater $K$ value than observed from 1000 samplings. Many of the approximated $K$ values are so close to 0 that the listed value is 0.

| Trait | Nuc | Treatment | Sex | $K$ | $p$ |
|---|---|---|---|---|---|
| Climbing | 375 | Control | F | 0.001 | 0.209 |
| Climbing | 375 | Control | M | 0.000 | 0.970 |
| Climbing | 375 | Rotenone | F | 0.000 | 0.940 |
| Climbing | 375 | Rotenone | M | 0.000 | 0.683 |
| Climbing | Ore | Control | F | 0.000 | 0.683 |
| Climbing | Ore | Control | M | 0.000 | 0.702 |
| Climbing | Ore | Rotenone | F | 0.001 | 0.180 |
| Climbing | Ore | Rotenone | M | 0.001 | 0.454 |
| Flight | 375 | Control | F | 0.000 | 0.980 |
| Flight | 375 | Control | M | 0.000 | 0.645 |
| Flight | 375 | Rotenone | F | 0.000 | 0.792 |
| Flight | 375 | Rotenone | M | 0.002 | 0.453 |
| Flight | Ore | Control | F | 0.000 | 0.841 |
| Flight | Ore | Control | M | 0.000 | 0.616 |
| Flight | Ore | Rotenone | F | 0.000 | 0.849 |
| Flight | Ore | Rotenone | M | 0.000 | 0.969 |
| Development | 375 | Control | MF | 0.000 | 0.718 |
| Development | 375 | Rotenone | MF | 0.000 | 0.650 |
| Development | Ore | Control | MF | 0.001 | 0.514 |
| Development | Ore | Rotenone | MF | 0.003 | 0.038 |
| Weight | 375 | Control | F | 0.000 | 0.864 |
| Weight | 375 | Control | M | 0.002 | 0.105 |
| Weight | 375 | Rotenone | F | 0.000 | 0.869 |
| Weight | 375 | Rotenone | M | 0.001 | 0.342 |
| Weight | Ore | Control | F | 0.001 | 0.382 |
| Weight | Ore | Control | M | 0.001 | 0.330 |
| Weight | Ore | Rotenone | F | 0.000 | 0.898 |
| Weight | Ore | Rotenone | M | 0.003 | 0.189 |

Supplementary Table 11: **All amino acid (AA) changes in mtDNA haplotypes.** Changes are given by complex, gene, and position (Pos.). Class is defined as radical when the AA transition is between groups (non-polar, uncharged polar, negatively charged, and positively charged) and conservative when the transition is within groups.

| Complex | Gene | Pos. | AA Change | Class. | Nuc. Contact | mtDNA |
|---|---|---|---|---|---|---|
| I | ND1 | 154 | L→M | Conservative | | yak |
| I | ND1 | 170 | F→Y | Radical | | sil |
| I | ND1 | 171 | F→Y | Radical | | yak |
| I | ND1 | 174 | V→I | Conservative | ✓ | yak |
| I | ND1 | 186 | A→S | Radical | ✓ | yak |
| I | ND1 | 190 | M→L | Conservative | ✓ | sil,yak |
| I | ND1 | 191 | S→T | Conservative | | sil,yak |
| I | ND1 | 258 | M→V | Conservative | | yak |
| I | ND1 | 270 | V→A | Conservative | | sil,yak |
| I | ND1 | 311 | L→F | Conservative | ✓ | yak |
| I | ND2 | 3 | N→Y | Conservative | ✓ | yak |
| I | ND2 | 11 | I→T | Radical | | yak |
| I | ND2 | 65 | V→A | Conservative | | sil,yak |
| I | ND2 | 81 | K→A | Radical | ✓ | yak |
| I | ND2 | 84 | M→L | Conservative | ✓ | sil,yak |
| I | ND2 | 148 | Y→F | Radical | | Zim53 |
| I | ND2 | 148 | Y→N | Conservative | | yak |
| I | ND2 | 192 | I→F | Conservative | ✓ | sil |
| I | ND2 | 202 | F→I | Conservative | | yak |
| I | ND2 | 239 | T→S | Conservative | ✓ | yak |
| I | ND2 | 268 | L→M | Conservative | ✓ | yak |
| I | ND2 | 274 | M→L | Conservative | | sil,yak |
| I | ND2 | 299 | M→L | Conservative | | yak |
| I | ND2 | 309 | K→E | Radical | ✓ | yak |
| I | ND2 | 311 | N→S | Conservative | ✓ | ZW140 |
| I | ND2 | 313 | N→I | Radical | ✓ | sil |
| I | ND2 | 315 | I→N | Radical | ✓ | sil,yak |
| I | ND2 | 317 | Y→T | Conservative | ✓ | sil,yak |
| I | ND2 | 319 | M→L | Conservative | | yak |
| I | ND2 | 321 | M→L | Conservative | ✓ | sil,yak |
| I | ND2 | 333 | L→M | Conservative | | sil |
| I | ND2 | 338 | Y→F | Radical | ✓ | yak |
| I | ND2 | 341 | F→L | Conservative | ✓ | yak |
| I | ND3 | 6 | F→I | Conservative | | yak |
| I | ND3 | 9 | L→S | Radical | | yak |
| I | ND3 | 10 | L→V | Conservative | | yak |
| I | ND3 | 17 | I→V | Conservative | | yak |
| I | ND3 | 20 | F→I | Conservative | | sil |
| I | ND3 | 29 | A→G | Radical | | sil |
| I | ND3 | 81 | M→I | Conservative | | sil |
| I | ND3 | 81 | M→L | Conservative | | yak |
| I | ND4 | 6 | F→L | Conservative | ✓ | yak |
| I | ND4 | 12 | I→T | Radical | | sil,yak |
| I | ND4 | 14 | F→L | Conservative | ✓ | sil |
| I | ND4 | 14 | F→V | Conservative | ✓ | yak |
| I | ND4 | 17 | I→M | Conservative | ✓ | sil |
| I | ND4 | 28 | M→L | Conservative | | yak |
| I | ND4 | 29 | F→V | Conservative | ✓ | sil |
| I | ND4 | 37 | L→V | Conservative | ✓ | sil |
| I | ND4 | 62 | I→V | Conservative | | yak |
| I | ND4 | 78 | M→S | Radical | | sil,yak |
| I | ND4 | 82 | H→Y | Radical | | sil,yak |

Continued on next page

Supplementary Table 11 – continued from previous page

| Complex | Gene | Pos. | AA Change | Class. | Nuc. Contact | mtDNA |
|---|---|---|---|---|---|---|
| I | ND4 | 84 | N→S | Conservative | ✓ | Bei05 |
| I | ND4 | 94 | I→V | Conservative | | sil,yak |
| I | ND4 | 101 | I→V | Conservative | | yak |
| I | ND4 | 141 | L→V | Conservative | | sil,yak |
| I | ND4 | 161 | L→V | Conservative | | ZW133, Bei04, Bei05, Bei11, Bei12, Bei23, Bei38, Bei42, Bei52, Bei59, ZH23, ZH26, ZH33, ZH42, ZW09, ZW140, ZW142, ZW144, sil, yak, Zim53 |
| I | ND4 | 165 | I→T | Radical | ✓ | yak |
| I | ND4 | 231 | M→L | Conservative | | sil,yak |
| I | ND4 | 236 | S→A | Radical | ✓ | sil |
| I | ND4 | 236 | S→N | Conservative | ✓ | yak |
| I | ND4 | 260 | V→M | Conservative | | yak |
| I | ND4 | 287 | S→A | Radical | | yak |
| I | ND4 | 298 | C→S | Conservative | | sil |
| I | ND4 | 342 | S→A | Radical | ✓ | yak |
| I | ND4 | 368 | Y→S | Conservative | | ZW133, Bei04, Bei05, Bei11, Bei12, Bei23, Bei38, Bei42, Bei52, Bei54, Bei59, ZH23, ZH26, ZH33, ZH42, ZW09, ZW140, ZW142, ZW144, sil, yak, Zim53 |
| I | ND4 | 383 | L→M | Conservative | ✓ | sil,yak |
| I | ND4 | 386 | F→L | Conservative | | sil |
| I | ND4 | 426 | L→F | Conservative | ✓ | sil |
| I | ND4 | 442 | F→C | Radical | ✓ | yak |
| I | ND4 | 443 | M→I | Conservative | | yak |
| I | ND4L | 57 | S→N | Conservative | | sil,yak |
| I | ND4L | 79 | V→I | Conservative | | sil |
| I | ND5 | 7 | V→I | Conservative | | sil,yak |
| I | ND5 | 8 | N→Y | Conservative | | sil |
| I | ND5 | 12 | M→I | Conservative | ✓ | sil,yak |
| I | ND5 | 15 | S→T | Conservative | | yak |
| I | ND5 | 23 | F→Y | Radical | | yak |
| I | ND5 | 27 | D→N | Radical | ✓ | sil,yak |
| I | ND5 | 29 | I→V | Conservative | ✓ | yak |
| I | ND5 | 36 | L→V | Conservative | ✓ | sil,yak |
| I | ND5 | 42 | M→S | Radical | ✓ | sil |
| I | ND5 | 65 | S→A | Radical | | sil,yak |
| I | ND5 | 77 | M→E | Radical | | yak |
| I | ND5 | 77 | M→S | Radical | | sil |
| I | ND5 | 78 | N→S | Conservative | | yak |
| I | ND5 | 80 | N→E | Radical | | yak |
| I | ND5 | 81 | H→N | Radical | | sil,yak |
| I | ND5 | 106 | I→V | Conservative | | yak |
| I | ND5 | 149 | L→F | Conservative | | sil |
| I | ND5 | 152 | S→A | Radical | | sil,yak |
| I | ND5 | 170 | I→V | Conservative | | yak |
| I | ND5 | 176 | E→S | Radical | ✓ | yak |
| I | ND5 | 184 | V→M | Conservative | | sil |
| I | ND5 | 234 | I→V | Conservative | | yak |
| I | ND5 | 244 | M→L | Conservative | | sil,yak |

Continued on next page

Supplementary Table 11 – continued from previous page

| Complex | Gene | Pos. | AA Change | Class. | Nuc. Contact | mtDNA |
|---|---|---|---|---|---|---|
| I | ND5 | 289 | L→Y | Radical | | yak |
| I | ND5 | 371 | V→I | Conservative | | yak |
| I | ND5 | 380 | Y→F | Radical | ✓ | yak |
| I | ND5 | 416 | I→V | Conservative | ✓ | yak |
| I | ND5 | 422 | M→L | Conservative | | sil,yak |
| I | ND5 | 426 | I→F | Conservative | | yak |
| I | ND5 | 450 | I→G | Radical | ✓ | yak |
| I | ND5 | 450 | I→T | Radical | ✓ | ZH26 |
| I | ND5 | 452 | M→L | Conservative | ✓ | yak |
| I | ND5 | 454 | L→M | Conservative | ✓ | yak |
| I | ND5 | 462 | V→M | Conservative | ✓ | sil |
| I | ND5 | 470 | I→V | Conservative | | sil |
| I | ND5 | 472 | L→I | Conservative | | yak |
| I | ND5 | 476 | F→Y | Radical | | sil,yak |
| I | ND5 | 477 | F→S | Radical | | yak |
| I | ND5 | 483 | F→L | Conservative | | yak |
| I | ND5 | 484 | M→N | Radical | ✓ | yak |
| I | ND5 | 486 | N→S | Conservative | ✓ | Bei04, Bei05, Bei11, Bei12, Bei23, Bei38, Bei52, Bei59 |
| I | ND5 | 488 | S→T | Conservative | | yak |
| I | ND5 | 489 | T→L | Radical | | yak |
| I | ND5 | 516 | V→I | Conservative | ✓ | sil |
| I | ND5 | 535 | Q→Y | Conservative | | yak |
| I | ND5 | 539 | M→N | Radical | ✓ | yak |
| I | ND5 | 539 | M→T | Radical | ✓ | sil |
| I | ND5 | 551 | S→F | Radical | ✓ | ZH26 |
| I | ND5 | 557 | L→M | Conservative | | sil,yak |
| I | ND5 | 564 | L→M | Conservative | | yak |
| I | ND5 | 567 | L→F | Conservative | | yak |
| I | ND5 | 567 | L→M | Conservative | | sil |
| I | ND5 | 569 | I→F | Conservative | | sil,yak |
| I | ND6 | 18 | L→F | Conservative | | yak |
| I | ND6 | 39 | L→M | Conservative | | sil |
| I | ND6 | 41 | T→S | Conservative | | yak |
| I | ND6 | 82 | M→I | Conservative | | sil,yak |
| I | ND6 | 86 | L→V | Conservative | | sil |
| I | ND6 | 89 | S→M | Radical | | yak |
| I | ND6 | 90 | L→F | Conservative | | yak |
| I | ND6 | 93 | I→F | Conservative | | yak |
| I | ND6 | 93 | I→L | Conservative | | sil |
| I | ND6 | 96 | L→F | Conservative | | yak |
| I | ND6 | 96 | L→M | Conservative | | sil |
| I | ND6 | 100 | F→L | Conservative | | sil |
| I | ND6 | 100 | F→M | Conservative | | yak |
| I | ND6 | 101 | I→V | Conservative | | sil |
| I | ND6 | 102 | M→I | Conservative | | sil |
| I | ND6 | 102 | M→L | Conservative | | yak |
| I | ND6 | 107 | S→F | Radical | | sil |
| I | ND6 | 107 | S→I | Radical | | yak |
| I | ND6 | 108 | S→T | Conservative | ✓ | yak |
| I | ND6 | 115 | D→E | Conservative | ✓ | yak |
| I | ND6 | 121 | N→E | Radical | | yak |
| I | ND6 | 127 | M→T | Radical | | yak |
| I | ND6 | 144 | I→V | Conservative | ✓ | yak |
| I | ND6 | 159 | I→V | Conservative | | sil,yak |

Continued on next page

Supplementary Table 11 – continued from previous page

| Complex | Gene | Pos. | AA Change | Class. | Nuc. Contact | mtDNA |
|---|---|---|---|---|---|---|
| III | Cyt-b | 2 | N→H | Radical | n/a | yak |
| III | Cyt-b | 60 | I→V | Conservative | n/a | yak |
| III | Cyt-b | 71 | C→Y | Conservative | n/a | ZH26 |
| III | Cyt-b | 99 | V→I | Conservative | n/a | yak |
| III | Cyt-b | 109 | K→L | Radical | n/a | yak |
| III | Cyt-b | 109 | K→M | Radical | n/a | sil |
| III | Cyt-b | 116 | I→V | Conservative | n/a | yak |
| III | Cyt-b | 264 | V→M | Conservative | n/a | Bei42 |
| III | Cyt-b | 324 | V→I | Conservative | n/a | yak |
| III | Cyt-b | 325 | M→L | Conservative | n/a | yak |
| III | Cyt-b | 357 | V→I | Conservative | n/a | sil,yak |
| III | Cyt-b | 358 | V→I | Conservative | n/a | yak |
| III | Cyt-b | 358 | V→M | Conservative | n/a | Bei04, Bei05, Bei11, Bei12, Bei23, Bei38, Bei52, Bei59 |
| III | Cyt-b | 365 | V→I | Conservative | n/a | yak |
| III | Cyt-b | 369 | I→V | Conservative | n/a | yak |
| IV | CoI | 2 | S→P | Radical | n/a | sil |
| IV | CoI | 106 | A→T | Radical | n/a | Bei52 |
| IV | CoI | 133 | A→S | Radical | n/a | yak |
| IV | CoI | 176 | S→T | Conservative | n/a | yak |
| IV | CoI | 412 | H→Q | Radical | n/a | sil,yak |
| IV | CoI | 451 | I→V | Conservative | n/a | yak |
| IV | CoI | 469 | F→Y | Radical | n/a | yak |
| IV | CoII | 115 | N→S | Conservative | n/a | sil |
| IV | CoII | 129 | M→A | Conservative | n/a | yak |
| IV | CoII | 129 | M→T | Radical | n/a | sil |
| IV | CoII | 130 | T→I | Radical | n/a | yak |
| IV | CoII | 143 | V→I | Conservative | n/a | sil,yak |
| IV | CoII | 218 | Y→H | Radical | n/a | sil |
| IV | CoII | 218 | Y→N | Conservative | n/a | yak |
| IV | CoIII | 41 | I→M | Conservative | n/a | sil |
| IV | CoIII | 45 | V→L | Conservative | n/a | sil,yak |
| IV | CoIII | 77 | T→I | Radical | n/a | Bei59 |
| IV | CoIII | 92 | V→I | Conservative | n/a | sil |
| IV | CoIII | 110 | A→S | Radical | n/a | ZW133 |
| IV | CoIII | 115 | A→T | Radical | n/a | Bei59 |
| IV | CoIII | 143 | V→I | Conservative | n/a | Bei04, Bei05, Bei11, Bei12, Bei23, Bei38, Bei52 |
| IV | CoIII | 155 | N→S | Conservative | n/a | yak |
| IV | CoIII | 170 | L→M | Conservative | n/a | Zim53 |
| IV | CoIII | 172 | I→V | Conservative | n/a | sil |
| IV | CoIII | 176 | I→M | Conservative | n/a | sil |
| IV | CoIII | 193 | I→V | Conservative | n/a | sil,yak |
| IV | CoIII | 199 | F→Y | Radical | n/a | sil,yak |
| IV | CoIII | 207 | I→V | Conservative | n/a | yak |
| IV | CoIII | 249 | V→I | Conservative | n/a | Bei04, Bei05, Bei11, Bei12, Bei23, Bei38, Bei52, Bei59 |
| V | ATPase6 | 17 | F→L | Conservative | n/a | sil,yak |
| V | ATPase6 | 28 | L→I | Conservative | n/a | sil |
| V | ATPase6 | 45 | M→F | Conservative | n/a | yak |
| V | ATPase6 | 45 | M→V | Conservative | n/a | sil |
| V | ATPase6 | 115 | L→M | Conservative | n/a | sil |

Continued on next page

Supplementary Table 11 – continued from previous page

| Complex | Gene | Pos. | AA Change | Class. | Nuc. Contact | mtDNA |
|---|---|---|---|---|---|---|
| V | ATPase6 | 120 | N→K | Radical | n/a | ZW133 |
| V | ATPase6 | 137 | I→V | Conservative | n/a | sil |
| V | ATPase6 | 185 | I→L | Conservative | n/a | sil,yak |
| V | ATPase6 | 185 | I→M | Conservative | n/a | Bei04, Bei05, Bei11, Bei12, Bei23, Bei38, Bei42, Bei52, Bei59, ZH33, ZW140, ZW142 |
| V | ATPase6 | 187 | V→I | Conservative | n/a | sil |
| V | ATPase6 | 192 | M→T | Radical | n/a | sil |
| V | ATPase6 | 192 | M→V | Conservative | n/a | yak |
| V | ATPase6 | 205 | A→T | Radical | n/a | yak |
| V | ATPase8 | 15 | I→V | Conservative | n/a | yak |
| V | ATPase8 | 26 | I→M | Conservative | n/a | sil |
| V | ATPase8 | 32 | M→I | Conservative | n/a | sil |
| V | ATPase8 | 34 | N→T | Conservative | n/a | yak |

Supplementary Table 12: **Amino acid changes in mtDNA genes.** Variant calls that map to amino acid (AA) changes for the *D. mel* haplotypes. Differences shown are between each sampled haplotype and the *D. mel* reference genome (REF). A dot indicates that there is no change. Radical changes between AA classes (non-polar, uncharged polar, negatively charged, and positively charged) are marked with $*$. Additionally, complex I site positions that contact nuclear-encoded proteins are marked with $*$. AA changes private to *D. sim* and/or *D. yak* are given in Table 11 and not shown here.

| | Complex I | | | | | | | | Complex III | | | Complex IV | | | | | | | Complex V | |
|---|---|---|---|---|---|---|---|---|---|---|---|---|---|---|---|---|---|---|---|---|
| | ND2 | | ND4 | | | ND5 | | | Cyt-b | | | CoI | CoIII | | | | | | ATPase6 | |
| | 148 | 311* | 84* | 161 | 368 | 450* | 486* | 551* | 71 | 264 | 358 | 106 | 77 | 110 | 115 | 143 | 170 | 249 | 120 | 185 |
| REF | Y | N | N | L | Y | I | N | S | C | V | V | A | T | A | A | V | L | V | N | I |
| Bei04 | . | . | . | V | S | . | S | . | . | . | M | . | . | . | . | I | . | I | . | M |
| Bei05 | . | . | S | V | S | . | S | . | . | . | M | . | . | . | . | I | . | I | . | M |
| Bei11 | . | . | . | V | S | . | S | . | . | . | M | . | . | . | . | I | . | I | . | M |
| Bei12 | . | . | . | V | S | . | S | . | . | . | M | . | . | . | . | I | . | I | . | M |
| Bei23 | . | . | . | V | S | . | S | . | . | . | M | . | . | . | . | I | . | I | . | M |
| Bei38 | . | . | . | V | S | . | S | . | . | . | M | . | . | . | . | I | . | I | . | M |
| Bei42 | . | . | . | V | S | . | . | . | . | M | . | . | . | . | . | . | . | . | . | M |
| Bei52 | . | . | . | V | S | . | S | . | . | . | M | T* | . | . | . | I | . | I | . | M |
| Bei54 | . | . | . | . | S | . | . | . | . | . | . | . | . | . | . | . | . | . | . | . |
| Bei59 | . | . | . | V | S | . | S | . | . | . | M | . | I* | . | T* | . | . | I | . | M |
| ZH23 | . | . | . | V | S | . | . | . | . | . | . | . | . | . | . | . | . | . | . | . |
| ZH26 | . | . | . | V | S | T* | . | F* | Y | . | . | . | . | . | . | . | . | . | . | . |
| ZH33 | . | . | . | V | S | . | . | . | . | . | . | . | . | . | . | . | . | . | . | M |
| ZH42 | . | . | . | V | S | . | . | . | . | . | . | . | . | . | . | . | . | . | . | . |
| ZW09 | . | . | . | V | S | . | . | . | . | . | . | . | . | . | . | . | . | . | . | . |
| ZW133 | . | . | . | V | S* | . | . | . | . | . | . | . | . | S | . | . | . | . | K* | . |
| ZW140 | . | S | . | V | S | . | . | . | . | . | . | . | . | . | . | . | . | . | . | M |
| ZW142 | . | . | . | V | S | . | . | . | . | . | . | . | . | . | . | . | . | . | . | M |
| ZW144 | . | . | . | V | S | . | . | . | . | . | . | . | . | . | . | . | . | . | . | . |
| Zim53 | F* | . | . | . | . | . | . | . | . | . | . | . | . | . | . | . | M | . | . | . |
| *D. sim* sil | . | . | . | V | S | . | . | . | . | . | . | . | . | . | . | . | . | . | . | . |
| *D. yak* | . | . | . | V | S | . | . | . | . | . | . | . | . | . | . | . | . | . | . | . |

Supplementary Table 14: **Weak and strong Mother's Curse hypothesis (MCH) tests for weight, flight, and climbing traits.** For each sex, the mean of the bootstrapped (n=1000) coefficient of variation (CV) distribution is given for females (Female Var.) and males (Male Var.). The $F$ value is and 95% confidence interval ($F_{lower}$,$F_{upper}$) are given for the $F$-ratio resulting from an $F$-test comparing male vs female variance, testing whether males have greater variance than females. A small $p$-value means males are significantly more variable than females. Sexual antagonism was tested by calculating the Pearson correlation coefficient $r$ between male and female performance, the value of $r$ and 95% confidence interval ($r_{lower}$,$r_{upper}$) are given. The $p$-value is given for a test with the alternative hypothesis that $r \neq 0$. The strong MCH necessitates a significant negative correlation. The Bonferroni corrected significance threshold is $\alpha = 0.05/12$ comparisons $= 0.004$.

| Trait | Nuc | Treatment | $F$ | $F_{lower}$ | $F_{upper}$ | $p$ |
|---|---|---|---|---|---|---|
| **Weak MCH** | | | | | | |
| weight | 375 | Control | 1.1715 | 0.4969 | 2.7624 | 0.3568 |
| weight | 375 | Rotenone | 0.5017 | 0.2083 | 1.2084 | 0.9391 |
| weight | Ore | Control | 0.5570 | 0.2362 | 1.3134 | 0.9111 |
| weight | Ore | Rotenone | 0.4371 | 0.1773 | 1.0771 | 0.9643 |
| flight | 375 | Control | 0.5399 | 0.2290 | 1.2729 | 0.9220 |
| flight | 375 | Rotenone | 3.0980 | 1.3139 | 7.3047 | 0.0053 |
| flight | Ore | Control | 0.9405 | 0.3989 | 2.2175 | 0.5566 |
| flight | Ore | Rotenone | 0.5312 | 0.2253 | 1.2524 | 0.9272 |
| climb | 375 | Control | 1.2219 | 0.5182 | 2.8810 | 0.3212 |
| climb | 375 | Rotenone | 1.1409 | 0.4839 | 2.6901 | 0.3800 |
| climb | Ore | Control | 0.7441 | 0.3156 | 1.7546 | 0.7530 |
| climb | Ore | Rotenone | 1.4610 | 0.6196 | 3.4448 | 0.1904 |
| | | | $r$ | $r_{lower}$ | $r_{upper}$ | $p$ |
| **Strong MCH** | | | | | | |
| weight | 375 | Control | 0.5402 | 0.1647 | 0.7790 | 0.0078 |
| weight | 375 | Rotenone | 0.7687 | 0.5136 | 0.8990 | 0.0000 |
| weight | Ore | Control | 0.4686 | 0.0699 | 0.7382 | 0.0241 |
| weight | Ore | Rotenone | 0.5584 | 0.1670 | 0.7978 | 0.0085 |
| flight | 375 | Control | 0.4549 | 0.0525 | 0.7302 | 0.0292 |
| flight | 375 | Rotenone | -0.2716 | -0.6149 | 0.1584 | 0.2100 |
| flight | Ore | Control | 0.6425 | 0.3132 | 0.8338 | 0.0009 |
| flight | Ore | Rotenone | 0.1216 | -0.3059 | 0.5084 | 0.5803 |
| climb | 375 | Control | 0.1007 | -0.3250 | 0.4925 | 0.6475 |
| climb | 375 | Rotenone | 0.4341 | 0.0267 | 0.7179 | 0.0385 |
| climb | Ore | Control | 0.2474 | -0.1835 | 0.5986 | 0.2550 |
| climb | Ore | Rotenone | 0.6637 | 0.3462 | 0.8448 | 0.0006 |

Supplementary Table 15: **dN/dS ratios for nucler-encoded OXPHOS genes.** This is a subset of the dN/dS ($\omega$) values calculated in Larracuente et. al [51] using *D. mel*, *D. sim*, and *D. yak*. The p-value ($p$) given is the for a PAML M7 vs M8 likelihood ratio test (LRT) with values <0.01 indicating positive selection for that gene. *D. mel* branch specific $\omega$ and $p$-values for positive selection are also given.

| Complex | Gene | $\omega$ | $p$ | *D. mel* $\omega$ | *D. mel* $p$ |
|---|---|---|---|---|---|
| I | ND-B17 | 0.04706 | 0.735428 | 0.0876 | 0.5000000 |
| I | ND-ACP | 0.03291 | 0.208389 | 0.0235 | 0.5000000 |
| I | ND-SGDH | 0.08386 | 0.735428 | 0.0001 | 0.5000000 |
| I | ND-23 | 0.01422 | 0.735428 | 0.0529 | 0.5000000 |
| I | ND-42 | 0.01562 | 0.283019 | 0.0001 | 0.5000000 |
| I | ND-PDSW | 0.05563 | 0.735428 | 0.1233 | 0.5000000 |
| I | ND-B14.5A | 0.02142 | 0.392941 | 0.0001 | 0.5000000 |
| I | NP15.6 | 0.03602 | 0.000084 | 0.0703 | 0.5000000 |
| I | ND-B16.6 | 0.04930 | 0.735428 | 0.1071 | 0.4666610 |
| I | ND-ASHI | 0.08551 | 0.069828 | 0.0001 | 0.5000000 |
| I | NdufV3 | 0.05406 | 0.023585 | 0.0001 | 0.5000000 |
| I | ND-20 | 0.05039 | 0.274259 | 0.0001 | 0.5000000 |
| I | ND-24 | 0.04756 | 0.735428 | 0.0904 | 0.5000000 |
| I | ND-18 | 0.04511 | 0.001937 | 0.0001 | 0.5000000 |
| I | ND-15 | 0.02487 | 0.735428 | 0.1207 | 0.4873864 |
| I | ND-B17.2 | 0.02207 | 0.735428 | 0.0663 | 0.5000000 |
| I | ND-B14.5B | 0.10698 | 0.001769 | 0.6241 | 0.1391881 |
| I | ND-13A | 0.08206 | 0.327746 | 0.0001 | 0.5000000 |
| I | ND-B22 | 0.04389 | 0.735428 | 0.0001 | 0.5000000 |
| I | ND-B15 | 0.05455 | 0.142015 | 0.1567 | 0.1154751 |
| I | ND-B14.7 | 0.02326 | 0.257075 | 0.0001 | 0.5000000 |
| I | ND-B12 | 0.04440 | 0.735428 | 0.1322 | 0.5000000 |
| I | ND-19 | 0.06949 | 0.358827 | 0.1022 | 0.5000000 |
| I | ND-B8 | 0.05063 | 0.735428 | 0.0397 | 0.5000000 |
| I | NdufA3 | 0.05375 | 0.360596 | 0.0001 | 0.5000000 |
| I | ND-13B | 0.06602 | 0.261203 | 0.0411 | 0.5000000 |
| II | SdhB | 0.00823 | 0.735428 | 0.0001 | 0.5000000 |
| II | SdhC | 0.02628 | 0.175539 | 0.0001 | 0.5000000 |
| II | SdhD | 0.06258 | 0.735428 | 0.2159 | 0.5000000 |
| III | ox | 0.04650 | 0.217234 | 0.0001 | 0.5000000 |
| III | RFeSP | 0.01408 | 0.023585 | 0.0001 | 0.5000000 |
| III | UQCR-6.4 | 0.10448 | 0.124579 | 0.0001 | 0.5000000 |
| III | Cyt-c1 | 0.04474 | 0.217571 | 0.0336 | 0.5000000 |
| III | UQCR-Q | 0.03861 | 0.735428 | 0.0001 | 0.5000000 |
| III | UQCR-C1 | 0.10338 | 0.211759 | 0.0001 | 0.5000000 |
| IV | COX5A | 0.01885 | 0.735428 | 999.0000 | 0.2572263 |
| IV | COX6B | 0.03826 | 0.408693 | 0.0709 | 0.5000000 |
| IV | COX5B | 0.03550 | 0.735428 | 0.1956 | 0.5000000 |
| IV | COX4 | 0.12349 | 0.407513 | 367.3428 | 0.5000000 |
| IV | levy | 0.04758 | 0.735428 | 0.0001 | 0.5000000 |
| IV | COX7A | 0.00916 | 0.735428 | 2.9970 | 0.5000000 |
| IV | COX7B | 0.03343 | 0.020637 | 0.0696 | 0.0986668 |
| IV | ND-MLRQ | 0.03752 | 0.148669 | 0.0001 | 0.5000000 |
| V | ATPsynG | 0.04577 | 0.402544 | 0.0646 | 0.4691289 |
| V | blw | 0.11224 | 0.735428 | 0.1683 | 0.5000000 |
| V | ATPsynCF6 | 0.03619 | 0.735428 | 0.0001 | 0.5000000 |
| V | ATPsynD | 0.04753 | 0.735428 | 0.0255 | 0.5000000 |
| V | ATPsynO | 0.01753 | 0.735428 | 0.0001 | 0.5000000 |
| V | ATPsynB | 0.01836 | 0.064185 | 0.0234 | 0.5000000 |
| V | ATPsyngamma | 0.07070 | 0.735428 | 0.0001 | 0.5000000 |
| V | ATPsyndelta | 0.01984 | 0.111860 | 0.0001 | 0.5000000 |

Continued on next page

Supplementary Table 15 – continued from previous page

| Complex | Gene | $\omega$ | $p$ | *D. mel* $\omega$ | *D. mel* $p$ |
|---|---|---|---|---|---|
| V | ATPsynF | 0.00526 | 0.348720 | 0.0001 | 0.5000000 |
| V | ATPsynC | 0.05204 | 0.280323 | 0.0001 | 0.5000000 |

Supplementary Table 16: **All mitonuclear genotypes tested.** For each mitonuclear genotype (*Mito;Nuc*) the exerimental block(s) where the line was tested is given. Both builds and all phenotypes were tested within each block unless specified otherwise. Mitochondrial haplogroups defined in Figure S2 in Early & Clark (2013) [70] are also given for each genotype. The flight trait was not measured in experimental block 9.

| Mitonuclear Genotype | Mitochondrial Haplogroup | Experimental Block(s) |
|---|---|---|
| *ZW144;OreR* | III | 1, 8 (development) |
| *ZW144;375* | III | 1, 8 (development) |
| *ZW142;OreR* | III | 1, 8 (development), 9 |
| *ZW142;375* | III | 1, 8 (development), 9 |
| *Bei59;OreR* | VIII | 1, 8 (development), 9 |
| *Bei59;375* | VIII | 1, 8 (development), 9 |
| *Bei54;OreR* | III | 1, 8 (development) |
| *Bei54;375* | III | 1, 8 (development) |
| *ZW133;OreR* | III | 2, 9 |
| *ZW133;375* | III | 2, 4 (build A), 9 |
| *ZH33;OreR* | III | 2, 9 |
| *ZH33;375* | III | 2, 9 |
| *Bei23;OreR* | VIII | 2, 9 |
| *Bei23;375* | VIII | 2, 9 |
| *Bei11;OreR* | VIII | 2, 9 |
| *Bei11;375* | VIII | 2, 9 |
| *ZW140;OreR* | I | 3 |
| *ZW140;375* | I | 3 |
| *ZW09;OreR* | III | 3 |
| *ZW09;375* | III | 3, 4 (build B) |
| *Bei52;OreR* | VIII | 3 |
| *Bei52;375* | VIII | 3 |
| *Bei42;OreR* | I | 3 |
| *Bei42;375* | I | 3 |
| *ZH42;OreR* | III | 4 |
| *ZH42;375* | III | 4 |
| *Bei05;OreR* | VIII | 4 |
| *Bei05;375* | VIII | 4 |
| *ZH26;OreR* | III | 5 |
| *ZH26;375* | III | 5 |
| *Bei04;OreR* | VIII | 5 |
| *Bei04;375* | VIII | 5 |
| *ZH23;OreR* | III | 6 |
| *ZH23;375* | III | 6 |
| *Bei38;OreR* | VIII | 6 |
| *Bei38;375* | VIII | 6 |
| *Zim53;OreR* | III | 7 |
| *Zim53;375* | III | 7 |
| *Bei12;OreR* | VIII | 7 |
| *Bei12;375* | VIII | 7 |
| *siI;OreR* | SIM | 1–3, 4–8 (build A), 9 |
| *siI;375* | SIM | 1–3, 4–8 (build A), 9 |
| *yak;OreR* | YAK | 1–3, 4–8 (build A), 9 |

Continued on next page

Supplementary Table 16 – continued from previous page

| Mitonuclear Genotype | Mitochondrial Haplogroup | Experimental Block(s) |
|---|---|---|
| *yak;375* | YAK | 1–3, 4–8 (build A), 9 |
| *OreR;OreR* | VI | 1–8 |
| *375;375* | VI | 1–8 |

### Supplementary References


[1] Bates, D., Mächler, M., Bolker, B. & Walker, S. Fitting linear mixed-effects models using lme4. *Journal of Statistical Software* **67**, 1–48 (2015).

[2] Kuznetsova, A., Brockhoff, P. B. & Christensen, R. H. B. lmerTest Package: tests in linear mixed effects models. *Journal of Statistical Software* **82**, 1–26 (2017).

[3] Lenth, R. V. *et al.* emmeans: Estimated marginal means, aka least-squares means (2025).

[4] Mendiburu, F. d. agricolae: Statistical procedures for agricultural research (2023).

[5] Li, H. & Durbin, R. Fast and accurate short read alignment with Burrows–Wheeler transform. *Bioinformatics* **25**, 1754–1760 (2009).

[6] Van der Auwera, G. A. *et al.* From FastQ data to high confidence variant calls: the Genome Analysis Toolkit best practices pipeline. *Current Protocols in Bioinformatics* **43**, 11.10.1–11.10.33 (2013).

[7] Cingolani, P. *et al.* A program for annotating and predicting the effects of single nucleotide polymorphisms, SnpEff: SNPs in the genome of Drosophila melanogaster strain $w^{1118}$ ; iso-2; iso-3. *Fly* **6**, 80–92 (2012).

[8] Schrödinger, LLC. 2015. The PyMOL Molecular Graphics System, Version 1.8. Unpublished.

[9] Katoh, K. & Standley, D. M. MAFFT multiple sequence alignment software version 7: improvements in performance and usability. *Molecular Biology and Evolution* **30**, 772–780 (2013).

[10] Kozlov, A.M., Darriba, D., Flouri, T., Morel, B., and Stamatakis, A. RAxML-NG: a fast, scalable and user-friendly tool for maximum likelihood phylogenetic inference. *Bioinformatics* **35**, 4453–4455 2019.

[11] Revell, L. J. phytools 2.0: an updated R ecosystem for phylogenetic comparative methods (and other things). *PeerJ* **12**, e16505 (2024).
